# A microtubule stability switch alters isolated vascular smooth muscle calcium flux in response to matrix rigidity

**DOI:** 10.1101/2022.11.23.517637

**Authors:** Robert Johnson, Finn Wostear, Reesha Solanki, Oliver Steward, Christopher Morris, Stefan Bidula, Derek T Warren

## Abstract

During ageing, the extracellular matrix of the aortic wall becomes more rigid. In response, VSMCs generate enhanced contractile forces. Our previous findings demonstrate that VSMC volume is enhanced in response to increased matrix rigidity, but our understanding of mechanisms regulating this process remain incomplete. In this current study, we show that microtubule stability in VSMCs is reduced in response to enhanced matrix rigidity via piezo1-mediated Ca^2+^ influx. Moreover, VSMC volume and Ca^2+^ flux was regulated by microtubule dynamics; microtubule stabilising agents reduced both VSMC volume and Ca^2+^ flux on rigid hydrogels, whereas microtubule destabilising agents increased VSMC volume and Ca^2+^ flux on pliable hydrogels. Finally, we show that disruption of the microtubule deacetylase HDAC6 uncoupled these processes and increased K40 alpha tubulin acetylation, VSMC volume and Ca^2+^ flux on pliable hydrogels, but did not alter VSMC microtubule stability. These findings uncover a microtubule stability switch that controls VSMC volume by regulating Ca^2+^ flux. Together, these data demonstrate that manipulation of microtubule stability can modify VSMC matrix rigidity response.

## Introduction

Maintaining aortic compliance, the ability of the aorta to change shape in response to changes in blood pressure is essential for cardiovascular (CV) health. Decreased aortic compliance is a major risk factor associated with a variety of age-related CV diseases (Glasser et al., 1997; Lacolley et al., 2020; Mitchell et al., 2010). The rigidity of the aortic wall is a major contributor to aortic compliance. In the healthy aortic wall, rigidity and compliance are determined by the balance between elastic-fibres, including elastin, that provide pliability, and non-elastic fibres, including collagen-I, that provide tensile strength, to the extracellular matrix (ECM) (Tsamis et al., 2013). However, during ageing and CV disease, elastic-fibres degrade, and collagen-I accumulates. These ECM changes increase the rigidity of the aortic wall and decrease aortic compliance (Afewerki et al., 2019; Ahmed and Warren, 2018; Johnson et al., 2021).

Vascular smooth muscle cells (VSMC) are the predominant cell type within the aortic wall. VSMC contraction regulates vascular tone in healthy aortae, where aortic wall rigidity and compliance are a balance between ECM rigidity and VSMC stiffness (Johnson et al., 2021). Changes in intracellular calcium ion (Ca^2+^) levels are integral for regulating VSMC contractile function (Ahmed and Warren, 2018). As such, intracellular Ca^2+^ are normally tightly regulated by mechanisms that promote release and reuptake of Ca^2+^ from intracellular stores such as the sarcoplasmic reticulum. During VSMC contraction, Ca^2+^ release is stimulated via receptor signalling or stretch activated ion channels, which include the TRPC family and piezo1, leading to the activation of sarcoplasmic IP_3_R and ryanodine receptors (Sanders, 2001). This results in a rapid Ca^2+^ spark in VSMCs that drives actomyosin force generation. These sparks are short lived as Ca^2+^ is rapidly reabsorbed to the sarcoplasmic reticulum via SERCA channels(Sanders, 2001).

However, the balance between ECM rigidity and VSMC stiffness is disrupted in ageing and CV disease, resulting in VSMC dysfunction (Lacolley et al., 2020; Tsamis et al., 2013). Age associated changes in ECM composition increase aortic wall rigidity and triggers increased VSMC stiffness, that further decreases aortic compliance (Lacolley et al., 2020; Sehgel et al., 2015, 2013; Tsamis et al., 2013).

This is termed VSMC stiffness syndrome, but our understanding of the mechanisms driving this process remain poorly defined. In response to hypertension, VSMCs undergo a process known as hypertrophy and increase their cell mass without increasing cell number (Owens and Schwartz, 1983; Rizzoni et al., 2000; Schiffrin, 2012; Zhang et al., 2005). Increases in volume and protein synthesis are key components in VSMC hypertrophy, which is known to increase aortic wall thickness and rigidity and decrease aortic compliance (Owens and Schwartz, 1983; Rizzoni et al., 2000; Schiffrin, 2012; Sehgel et al., 2013; Zhang et al., 2005). Importantly, we have recently shown that activation of piezo1 drives a sustained Ca^2+^ influx, resulting in an increase in VSMC volume, via the membrane translocation of the water channel aquaporin-1, in response to matrix rigidity (Johnson et al., 2024). Increased matrix rigidity and hypertension are known to promote VSMC dedifferentiation, where they switch from a contractile phenotype to disease associated synthetic phenotypes (Cao et al., 2022; Hayashi and Naiki, 2009; Xie et al., 2018). Both aquaporin-1 and piezo1 gene expression is enhanced during VSMC dedifferentiation in atherosclerosis, during carotid remodelling and during aortic ageing (Johnson et al., 2024; Luu et al., 2023). Whether increased volume drives increased VSMC stiffness in response to matrix rigidity and the VSMC phenotypes involved remain unknown.

Microtubules are hollow tube-like structures, made up of alpha and beta tubulin heterodimers, that play important roles in determining cell morphology and resisting deformational forces applied to cells (Brangwynne et al., 2006; Stamenović, 2005). Microtubule organisation is regulated via a process called dynamic instability, where microtubules exist in a balance between growth and shrinkage that enables them to respond rapidly to the changing needs of a cell (Michaels et al., 2020). Post-translational modifications of tubulin are predicted to regulate microtubule mechanical properties (Song and Brady, 2015). Most modifications are predicted to occur on the outer surface of polymerised microtubules. However, lysine 40 (K40) of alpha-tubulin in the inner lumen of polymerised microtubules, can be acetylated and deacetylated by αTAT1 and HDAC6, respectively (Osseni et al., 2020; Shida et al., 2010). While the precise impact of K40 acetylation on microtubule mechanical properties remains controversial and contradictory, K40 acetylation is proposed to influence lateral associations between polymerised tubulin monomers and potentially stabilise microtubules, making them more resistant to mechanical damage than deacetylated microtubules (Eshun-Wilson et al., 2019; Janke and Montagnac, 2017; Xu et al., 2017). Enhanced K40 acetylation increases cytoskeletal stiffness in striated muscle, but the impact of K40 acetylation in VSMC matrix rigidity sensing remains unknown(Coleman et al., 2021).

The mechanical regulation of VSMC behaviour obeys well defined rules. For example, in response to enhanced matrix rigidity, VSMCs generate increased actomyosin derived traction forces and increased deformational stresses are placed upon the cell membrane (Petit et al., 2019; Sanyour et al., 2019). These deformational stresses drive the opening of stretch-activated ion channels in VSMCs (Johnson et al., 2024). Microtubules exist in a mechanical balance with actomyosin activity, serving as pre-stressed compression bearing struts capable of resisting actomyosin generated deformational stresses (Brangwynne et al., 2006; Johnson et al., 2021; Stamenović, 2005). This relationship is known as the tensegrity model (Stamenović, 2005). The tensegrity model predicts the interplay between actomyosin and microtubules in VSMCs. Previous studies have shown that microtubule stabilisation decreases VSMC actomyosin activity, whereas microtubule destabilisation increases VSMC actomyosin activity (Ahmed et al., 2021; Zhang et al., 2000). Other models have been proposed that describe VSMC mechano-adaption responses, yet we lack an in-depth mechanistic understanding of pathways linking changes in VSMC behaviour to the proposed models of mechanotransduction (Steucke et al., 2017). For example, whether mechanical cues and stretch-activated ion channel activity influence microtubule stability and vice versa remains unknown.

In this current study, we show that changes in microtubule stability serve as a switch that regulates Ca^2+^ flux and VSMC volume control. Treatment with the microtubule stabilising agent paclitaxel reduced VSMC volume on rigid hydrogels. In contrast, treatment with the microtubule destabilising agent colchicine increased VSMC volume on pliable hydrogels. Furthermore, we show that paclitaxel reduced Ca^2+^ flux specifically in VSMCs on rigid hydrogels, whereas colchicine treatment increased Ca^2+^ flux in VSMCs on pliable hydrogels. In keeping with our previous findings, our data also implicates piezo1 in this process and depletion of piezo1 increased microtubule stability specifically on rigid hydrogels. Finally, we demonstrate that disruption of HDAC6 increases acetylation of alpha tubulin and induces increased volume and Ca^2+^ flux in VSMCs on pliable hydrogels. Together, these data implicate a piezo1/Ca^2+/^microtubule stability feedback pathway in the regulation of VSMC matrix rigidity response.

## Results

### Piezo1 activity promotes microtubule destabilisation within VSMCs on rigid hydrogels

Previous studies have shown that matrix rigidity promotes increased VSMC traction stress generation. To confirm that angiotensin II stimulated VSMCs seeded on rigid hydrogels generated enhanced traction stresses, we performed traction force microscopy. As expected, analysis confirmed that angiotensin II stimulated VSMCs seeded on rigid hydrogels generated greater maximal and total traction stress, compared to their counterparts on pliable hydrogels (**Supplementary Figure S1**). Our previous study found that on pliable hydrogels, VSMCs generated enhanced traction stresses as a result of colchicine-induced microtubule depolymerisation. Therefore, we predicted that VSMCs would display decreased microtubule stability on rigid hydrogels. To test this, we compared the number of cold-stable microtubules possessed by quiescent and angiotensin II stimulated VSMCs on pliable and rigid hydrogels. Analysis revealed that angiotensin II stimulation resulted in a reduction of cold-stable microtubules possessed by VSMCs on pliable and rigid hydrogels (**Figure 1 A and B**). However, this decrease was only significantly different on rigid hydrogels (**Figure 1A and B**).

**Figure 1.**
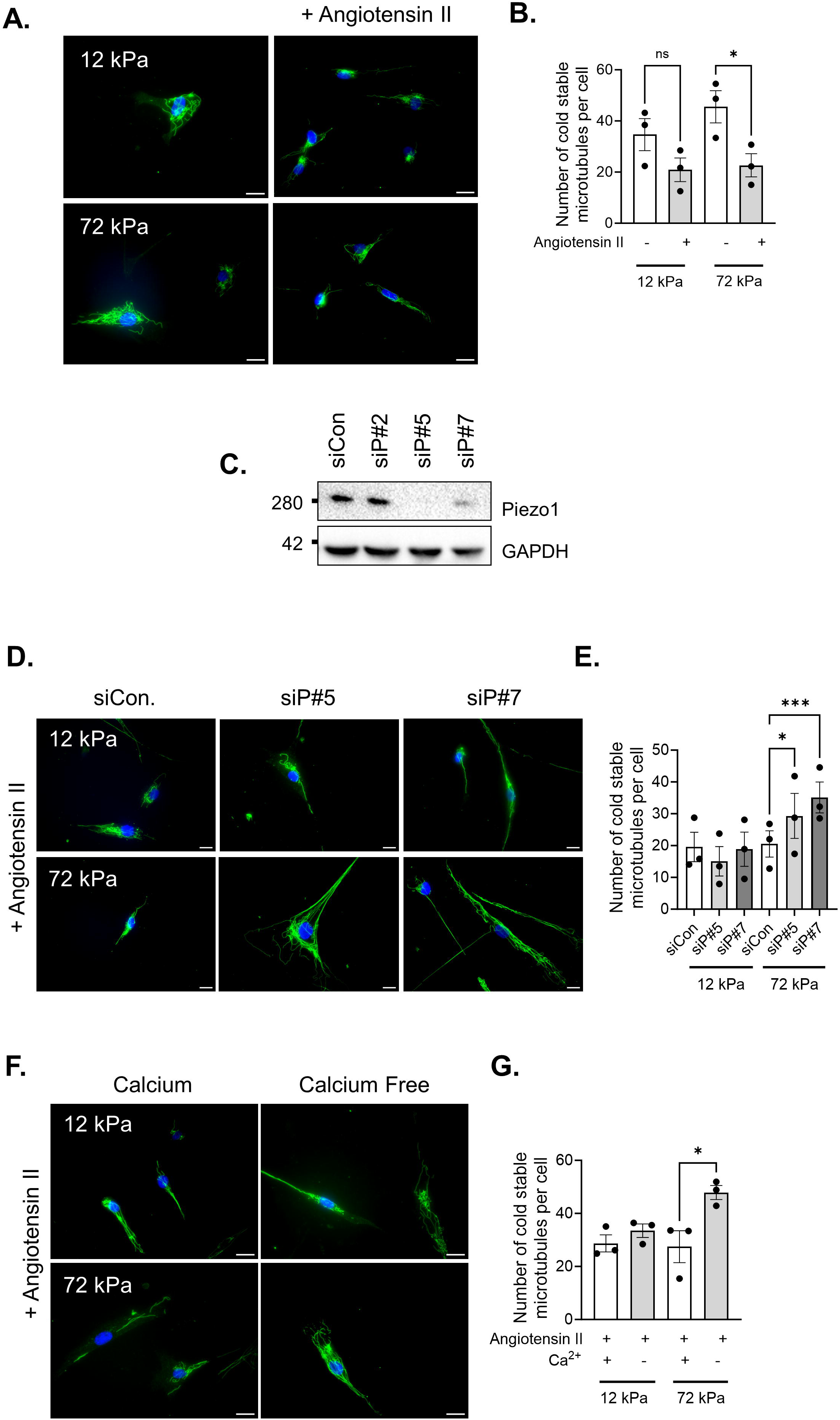
Extracellular calcium influx via piezo1 ion channels triggers microtubule destabilisation. A) Representative images of isolated VSMCs cultured on 12 or 72 kPa polyacrylamide hydrogels with/without angiotensin II treatment. Cold-stable microtubules, (α-tubulin, green) and nuclei (DAPI, blue). Scale bar = 50 μm. B) Graph shows number of cold-stable microtubules per cell on pliable and rigid hydrogels with/without angiotensin II treatment. Graph represents combined data from 3-independent experiments, analysing >56 cells per condition, and black dots mark mean data for each independent repeat. C) Representative WB confirming efficient siRNA-mediated piezo1 depletion in VSMCs (siCon = non-targeting scrambled siRNA, siP#5 and siP#7 = independent piezo1 targeting siRNA #5 and #7). D) Representative images of angiotensin II stimulated siCon, siP#5 and siP#7 treated VSMCs cultured on 12 or 72 kPa polyacrylamide hydrogels. Cold-stable microtubules, (α-tubulin, green) and nuclei (DAPI, blue). Scale bar = 50 μm. E) Graph shows number of cold-stable microtubules per cell on 12 kPa and 72 kPa hydrogels in angiotensin II stimulated siCon, siP#5 and siP#7 VSMCs. Graph represents combined data from 3-independent experiments, analysing >65 cells per condition, and black dots mark mean data for each independent repeat. F) Representative images of angiotensin II stimulated VSMCs cultured on 12 or 72 kPa polyacrylamide hydrogels in calcium containing or calcium free media. Cold-stable microtubules, (α-tubulin, green) and nuclei (DAPI, blue). Scale bar = 50 μm. G) Graph shows number of cold-stable microtubules per cell on 12 kPa and 72 kPa hydrogels in angiotensin II stimulated VSMCs in the presence/absence of extracellular calcium. Graph represents combined data from 3-independent experiments, analysing >71 cells per condition, and black dots mark mean data for each independent experimental repeat. Significance determined using a two-way ANOVA followed by Tukey’s post-hoc test. (* = p < 0.05, ** = p < 0.01 and *** = p < 0.001. Error bars represent ± SEM)

We have previously shown that piezo1 is an essential mediator of the enhanced VSMC volume response on rigid hydrogels. We next utilised a siRNA mediated knockdown approach to investigate whether piezo1 was required for this change in microtubule stability. Consistent with our previous studies, western blot analysis confirmed that piezo1 was efficiently depleted in VSMCs by piezo1 specific siRNA compared to non-targeting scrambled siRNA (**Figure 1C**). Analysis of microtubule stability revealed that piezo1 depleted, angiotensin II stimulated VSMCs possessed similar numbers of cold stable microtubules on pliable hydrogels as their scrambled control counterparts (**Figure 1 D and E**). In contrast, piezo1 depleted angiotensin II stimulated VSMCs on rigid hydrogels possessed increased numbers of cold-stable microtubules compared to their scrambled control counterparts (**Figure D and E**). We have previously shown that piezo1 activation promotes Ca^2+^ influx in VSMCs on rigid hydrogels. To confirm that Ca^2+^ influx was driving microtubule destabilisation, we performed a cold-stable microtubule assay in the presence or absence of extracellular Ca^2+^. The absence of extracellular Ca^2+^ had no effect on the number of cold-stable microtubules detected in angiotensin II stimulated VSMCs on pliable hydrogels (**Figure 1F and G**). In contrast, the number of cold-stable microtubules increased within angiotensin II stimulated VSMCs on rigid hydrogels when extracellular Ca^2+^ was absent (**Figure 1F and G**). This suggests that piezo1-mediated Ca^2+^ influx decreases microtubule stability on rigid hydrogels following angiotensin II stimulation.

### Microtubule stability influences isolated smooth muscle cell volume following contractile agonist stimulation

The above data shows that on rigid substrates, VSMCs generate increased traction stresses and possess decreased microtubule stability. We have previously shown that VSMCs on rigid hydrogels swell and possess increased cell area and volume. We next hypothesised that decreased microtubule stability was driving the enhanced VSMC volume response on rigid hydrogels. To test this, we utilised our screening assay described previously. Quiescent VSMCs were seeded onto pliable and rigid hydrogels and pre-treated with increasing concentrations of microtubule stabilisers prior to angiotensin II stimulation. Treatment with either paclitaxel or epothilone B had no effect on the contractile response of VSMCs seeded on pliable hydrogels (**Figure 2A, B, D and Supplementary Figure S2**). In contrast, increasing concentrations of either microtubule stabiliser was sufficient to prevent the increase in VSMC area observed following angiotensin II stimulation on rigid hydrogels (**Figure 2 A, C, D and Supplementary Figure S2**). Given that microtubule stabilisation prevented VSMC enlargement following angiotensin II stimulation on rigid hydrogels, we next hypothesised that microtubule destabilisation would trigger VSMC enlargement in angiotensin II stimulated VSMCs on pliable hydrogels.

**Figure 2.**
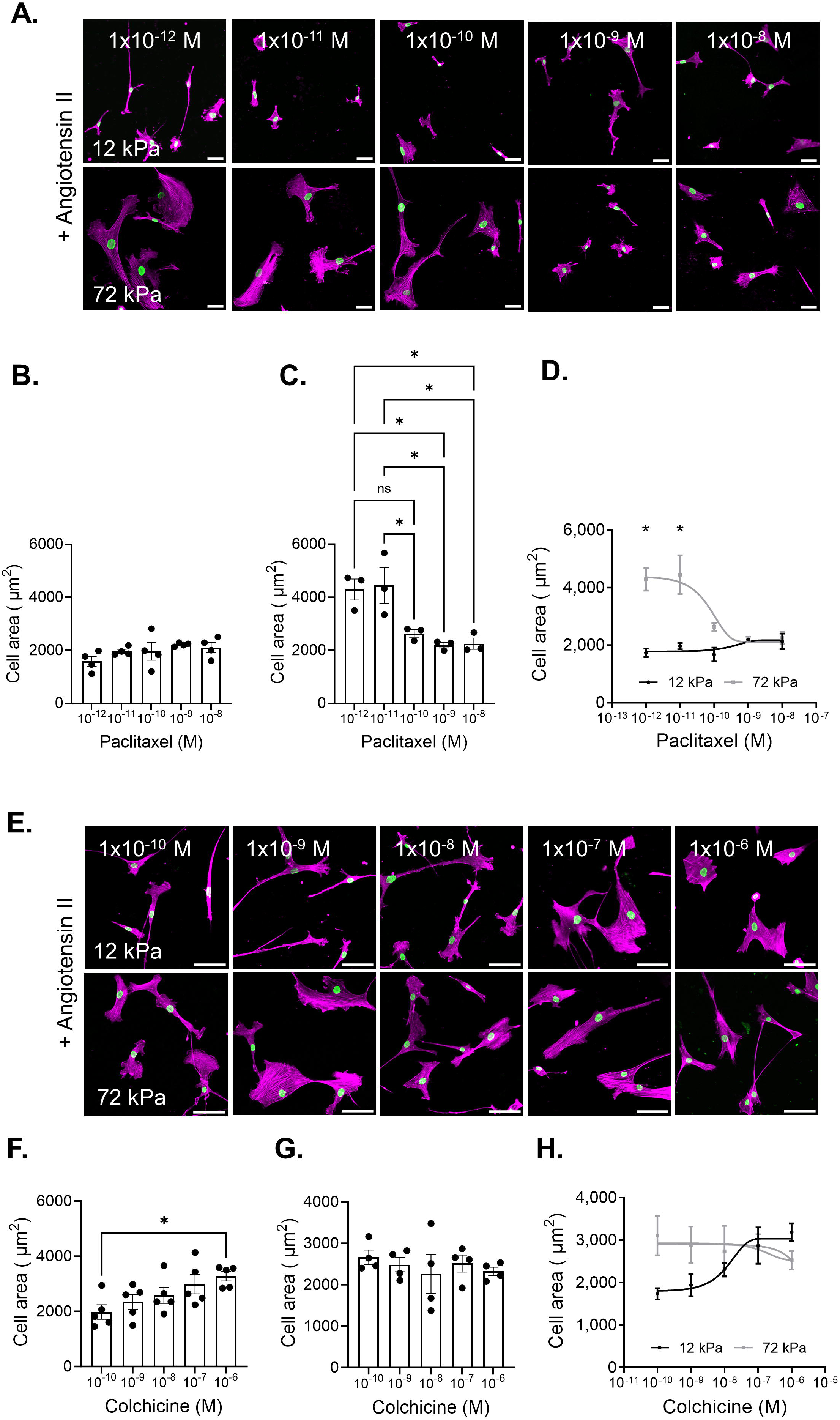
Microtubule stability influences VSMC spreading following angiotensin II stimulation. A) Representative images of isolated VSMCs cultured on 12 or 72 kPa polyacrylamide hydrogels pretreated with a concentration range of paclitaxel prior to angiotensin II stimulation. Actin cytoskeleton (purple) and lamin A/C labelled nuclei (green). Scale bar = 50 μm. Graphs show VSMC area on B) 12 kPa hydrogels, C) 72 kPa hydrogels, and D) 12 kPa and 72 kPa hydrogels. Graphs represent the combined data from 3-independent experiments, analysing <80 cells per condition. Black dots represent mean data from each individual experimental repeat. E) Representative images of isolated VSMCs cultured on 12 or 72 kPa polyacrylamide hydrogels pretreated with a concentration range of colchicine prior to angiotensin II stimulation. Actin cytoskeleton (purple) and lamin A/C labelled nuclei (green). Scale bar = 50 μm. Graphs show VSMC area on F) 12 kPa hydrogels, G) 72 kPa hydrogels, and H) 12 kPa and 72 kPa hydrogels. Graphs represent the combined data from 3-independent experiments, analysing <80 cells per condition. Black dots represent mean data from each individual experimental repeat. Significance determined using one-way ANOVA (graphs B, C, F and G) or two-way ANOVA (Graphs D and H) followed by Tukey’s post-hoc test. (* = p < 0.05. Error bars represent ± SEM)

VSMCs were pre-treated with increasing concentrations of the microtubule destabilisers colchicine or nocodazole, and then stimulated with angiotensin II. On pliable hydrogels, VSMCs treated with either colchicine or nocodazole displayed increased cell area following angiotensin II stimulation (**Figure 2E, F, H and Supplementary Figure S3**). In contrast, treatment with the microtubule destabilisers had no effect on VSMCs seeded on rigid hydrogels (**Figure 2E, G, H and Supplementary Figure S3**). All microtubule targeting agents were used at concentrations that did not cause cell death, as confirmed through a viability assay for concentrations of epothilone B and nocodazole (**Supplementary Figure S4**) or previously for paclitaxel and colchicine (Ahmed et al., 2022). We next confirmed that paclitaxel and colchicine treatments were inducing microtubule stabilisation and destabilisation respectively. Quiescent VSMCs seeded on pliable and rigid hydrogels were pretreated with microtubule targeting agents prior to angiotensin II stimulation. Analysis confirmed that paclitaxel treatment (1 nM) resulted in increased numbers of cold stable microtubules in angiotensin II stimulated VSMCs on both rigid and pliable hydrogels (**Supplementary Figure S5**). Colchicine treatment (100 nM) resulted in reduced numbers of cold stable microtubules in angiotensin II stimulated VSMCs on pliable hydrogels but did not significantly alter the number of cold stable microtubules in angiotensin II treated VSMCs on rigid hydrogels (**Supplementary Figure S5**).

Having determined that microtubule stability regulated changes in VSMC area, we then sought to confirm its regulation of VSMC volume. Quiescent VSMCs were pre-treated with paclitaxel (1 nM) or colchicine (100 nM), prior to angiotensin II stimulation and confocal microscopy was used to assess changes in cell volume.

Treatment with the microtubule stabiliser, paclitaxel had no effect on the volume of VSMCs seeded on pliable hydrogels (**Figure 3A-D**). Whereas, on rigid hydrogels, microtubule stabilisation prevented the angiotensin II induced expansion of VSMC volume (**Figure 3A-D**). Finally, microtubule destabilisation, via colchicine pre-treatment, increased VSMC volume on pliable hydrogels following angiotensin II stimulation (**Figure 3D-F**) but had no additional effect on VSMC volume on rigid hydrogels (**Figure 3D-F**).

**Figure 3.**
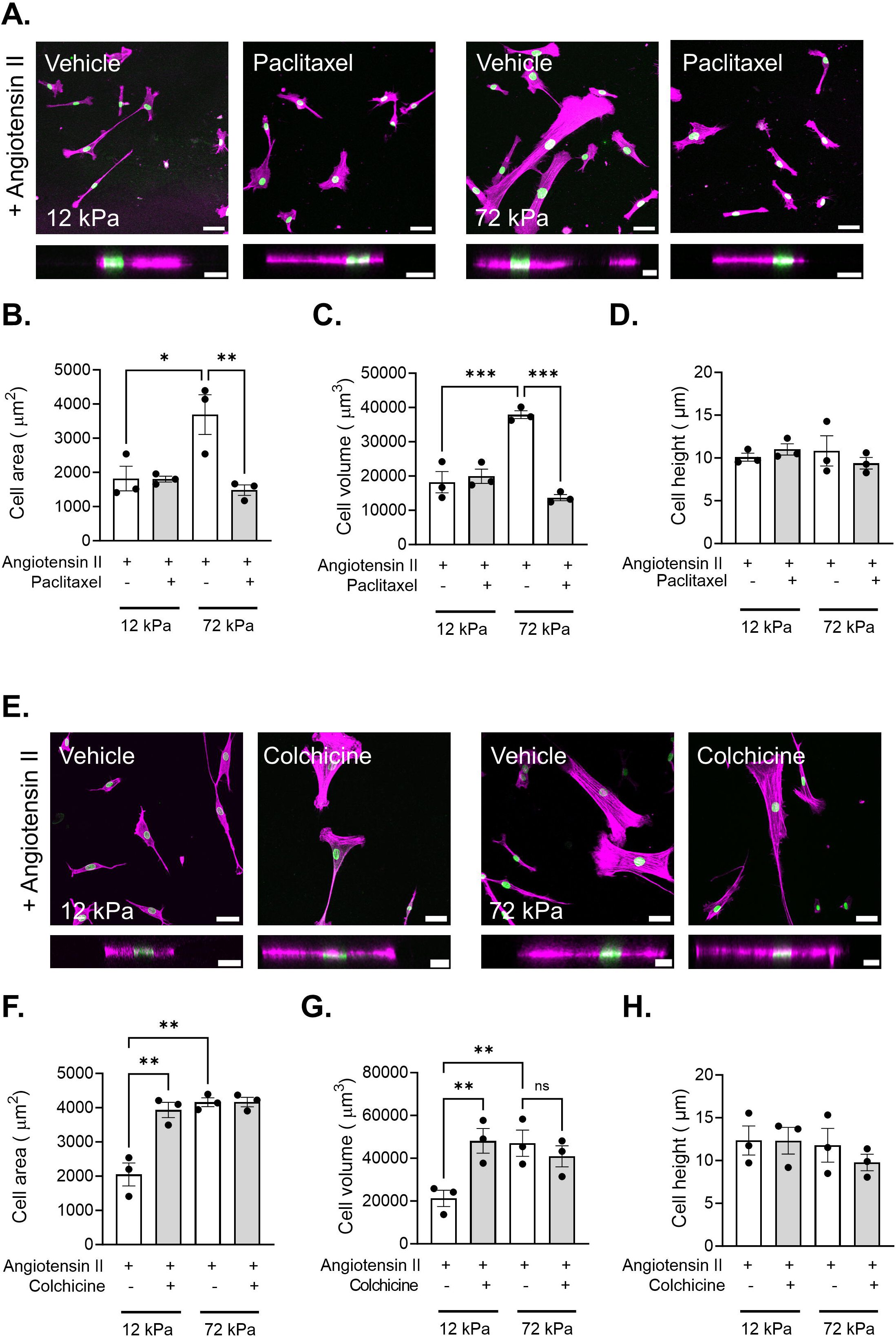
VSMC volume is regulated by microtubule stability. A) Representative images of isolated VSMCs cultured on 12 or 72 kPa polyacrylamide hydrogels pretreated with vehicle control (dH_2_0) or paclitaxel prior to angiotensin II stimulation. Actin cytoskeleton (purple) and DAPI labelled nuclei (green). Top panel shows representative XY images of VSMC area, scale bar = 50 μm. Bottom panel shows representative XZ images of VSMC height, scale bar = 20 μm. Graphs show VSMC B) area, C) volume and D) height and represent the combined data from 3 independent experiments with ≥45 cells analysed per condition. Mean data from each individual repeat of the 3 independent experiments is shown by a black dot. Significance was determined using two-way ANOVA followed by Tukey’s post-hoc test. E) Representative images of isolated VSMCs cultured on 12 or 72 kPa polyacrylamide hydrogels pretreated with vehicle control (dH_2_0) or colchicine prior to angiotensin II stimulation. Actin cytoskeleton (purple) and DAPI labelled nuclei (green). Top panel shows representative XY images of VSMC area, scale bar = 50 μm. Bottom panel shows representative XZ images of VSMC height, scale bar = 20 μm. Graphs show VSMC F) area, G) volume and H) height and represent the combined data from 3 independent experiments with ≥42 cells analysed per condition. Mean data from each individual repeat of the 3 independent experiments is shown by a black dot. Significance was determined using two-way ANOVA followed by Tukey’s test. (* = p < 0.05, ** = p < 0.01 and *** = p < 0.001. Error bars represent ± SEM)

### Changes in microtubule stability alter calcium ion flux

The above data confirm that microtubule stability influences VSMC volume control. Previously, we have identified piezo1 mediated Ca^2+^ influx as a regulator of VSMC volume control. Therefore, we next hypothesised that microtubule stability was altering Ca^2+^ flux in VSMCs. To test this, quiescent VSMCs on rigid and pliable hydrogels were loaded with the Ca^2+^ indicator Fluo-4 prior to angiotensin II treatment. Fluorescence video time lapse microscopy was used to measure changes in Fluo-4 fluorescence. Analysis supported our previous findings showing that Ca^2+^ flux was heightened and prolonged in VSMCs on rigid hydrogels compared to VSMCs on pliable hydrogels (**Figure 4A and B**). Colchicine treatment resulted in prolonged Ca^2+^ flux in VSMCs on pliable hydrogels but had little effect on Ca^2+^ flux in VSMCs on rigid hydrogels (**Figure 4A and B**). In contrast, paclitaxel treatment reduced the Ca^2+^ flux in VSMCs on rigid hydrogels but had no effect on Ca^2+^ flux in VSMCs on pliable hydrogels (**Figure 4A and B**).

**Figure 4.**
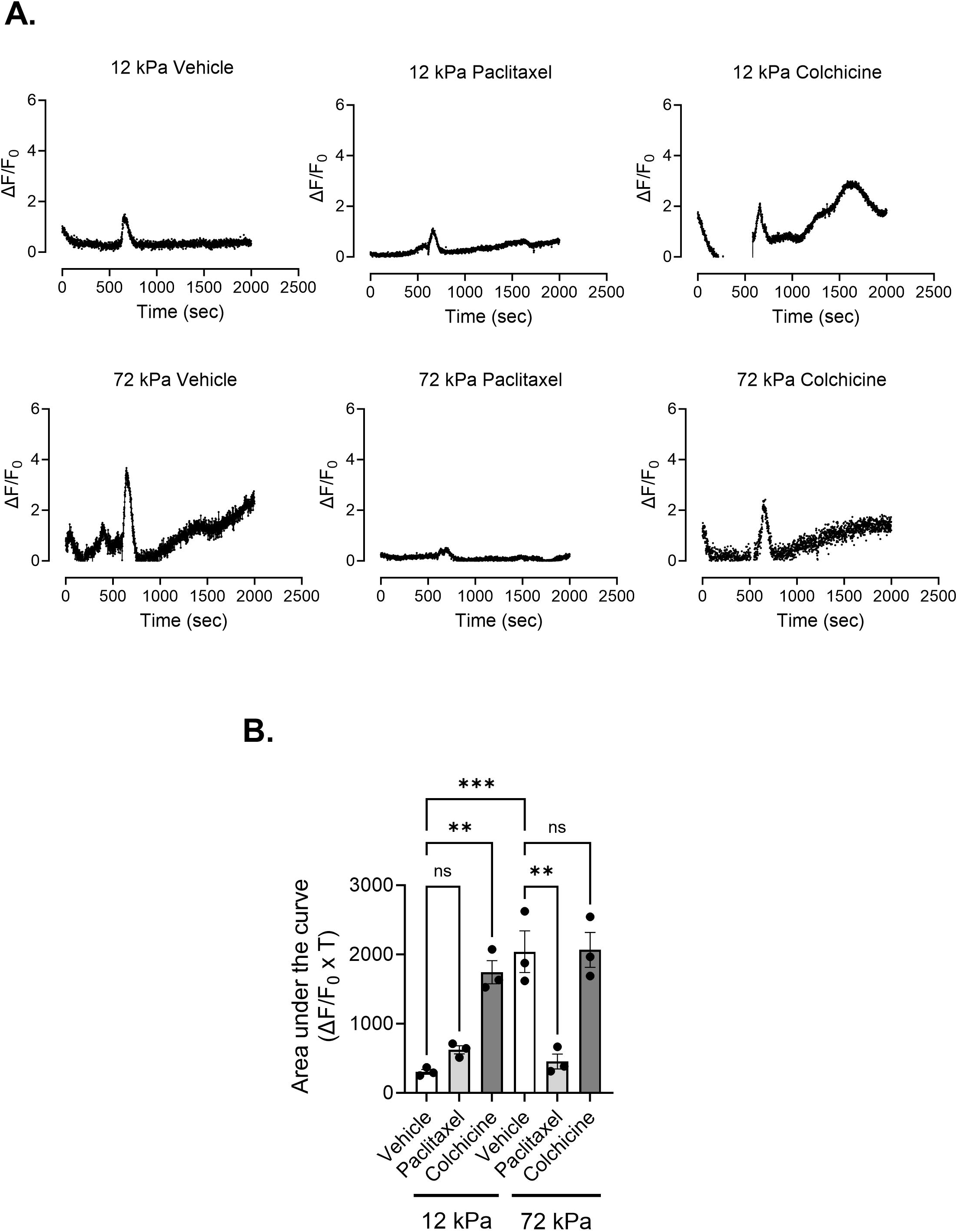
Microtubule stability influences Ca^2+^ flux in VSMCs. A) Graphs show representative ΔF/F_0_ values over time for vehicle control, paclitaxel and colchicine pre-treated angiotensin II stimulated Fluo-4 loaded VMSCs on 12 and 72 kPa hydrogels. Each graph represents the mean combined data of 9 individual fields of view combined from 3 individual experiments. B) Graph shows area under the curve values (ΔF/F_0_ x T). Mean data for individual repeats of the 3 independent experiments are shown as black dots. Significance was determined using two-way ANOVA followed by Tukey’s multiple comparison test. (** = p < 0.01 and *** = p < 0.001. Error bars represent ± SEM)

### HDAC6 disruption increases microtubule acetylation and induces enhanced VSMC volume response on pliable hydrogels

Although controversial, alpha tubulin K40 acetylation is proposed to alter the mechanical properties of microtubules. We next hypothesised that increased tubulin acetylation would influence the VSMC matrix rigidity response. To test this, we utilised tubastatin to inhibit HDAC6 deacetylation of the K40 position. Western blot analysis confirmed that VSMCs exposed to an increasing concentration of tubastatin possessed increased levels of acetylated alpha tubulin (**Supplementary Figure S6A and B**). Next, quiescent VSMCs were pretreated with tubastatin prior to angiotensin II stimulation and confocal microscopy was used to assess changes in the cell volume. Tubastatin treated VSMCs possessed increased volume compared to their vehicle treated counterparts on pliable hydrogels (**Figure 5A-D**). Tubastatin treatment had no effect on VSMC volume on rigid hydrogels (**Figure 5A-D**). To confirm that this altered response was driven by HDAC6 disruption, we performed siRNA mediated HDAC6 depletion experiments. WB confirmed HDAC6 was efficiently depleted in VSMCs by 2 independent siRNA oligomers (**Supplementary Figure S6C and D**). Analysis of z-stacks captured by confocal microscopy confirmed that HDAC6 depleted, angiotensin II stimulated VSMCs possessed increased volume on pliable hydrogels, compared to their scrambled control treated counterparts (**Figure 5E-H**). HDAC6 knockdown had no effect on VSMC volume on rigid hydrogels (**Figure 5E-H**). Due to the increased volume of HDAC6 disrupted VSMCs on pliable hydrogels, we next predicted that increased tubulin acetylation was altering VSMC microtubule stability and Ca^2+^ flux. To test these possibilities, we firstly performed cold-stable microtubule stability assays in tubastatin treated VSMCs on pliable and rigid hydrogels. However, the number of cold-stable microtubules remained unaffected by tubastatin treatment (**Figure 6A and B**). Secondly, quiescent VSMCs on rigid and pliable hydrogels were loaded with the Ca^2+^ indicator Fluo-4 prior to angiotensin II treatment. Fluorescence video time lapse microscopy was used to measure changes in Fluo-4 fluorescence. Tubastatin treatment resulted in prolonged Ca^2+^ flux in VSMCs on pliable hydrogels but had little effect on Ca^2+^ flux in VSMCs on rigid hydrogels (**Figure 6E and F**). Finally, we investigated whether matrix rigidity influenced acetylated K40 α-tubulin levels. However, WB analysis revealed that acetylated K40 levels were similar in the lysates of VSMCs grown on 12 and 72 kPa hydrogels (**Supplementary Figure S7**).

**Figure 5.**
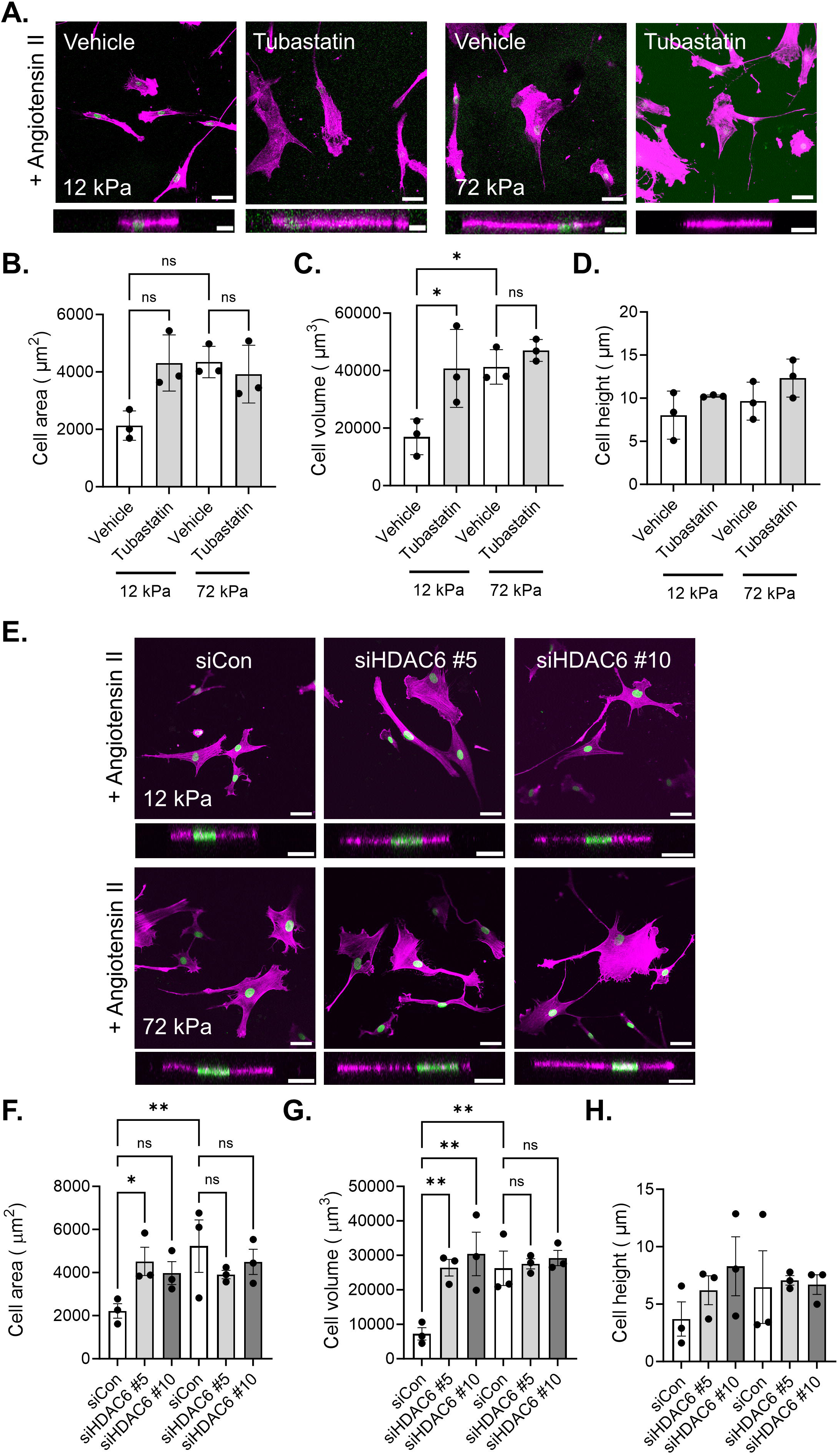
HDAC6 disruption increases microtubule acetylation and enhances VSMC volume. A) Representative images of isolated VSMCs cultured on 12 or 72 kPa polyacrylamide hydrogels pretreated with vehicle control (dH_2_0) or tubastatin prior to angiotensin II stimulation. Actin cytoskeleton (purple) and DAPI labelled nuclei (green). Top panel shows representative XY images of VSMC area, scale bar = 50 μm. Bottom panel shows representative XZ images of VSMC height, scale bar = 20 μm. Graphs show VSMC B) area, C) volume and D) height and represent the combined data from 3 independent experiments with ≥41 cells analysed per condition. Mean data from each individual repeat of the 3 independent experiments is shown by a black dot. Significance was determined using two-way ANOVA followed by Tukey’s test. E) Representative images of isolated VSMCs cultured on 12 or 72 kPa polyacrylamide hydrogels treated with scrambled (siCon) or HDAC6 targeting (HDAC6 #5 and #10) siRNA oligos prior to angiotensin II stimulation. Actin cytoskeleton (purple) and DAPI labelled nuclei (green). Top panel shows representative XY images of VSMC area, scale bar = 50 μm. Bottom panel shows representative XZ images of VSMC height, scale bar = 20 μm. Graphs show VSMC F) area, G) volume and H) height and represent the combined data from 3 independent experiments with ≥30 cells analysed per condition. Mean data from each individual repeat of the 3 independent experiments is shown by a black dot. Significance was determined using two-way ANOVA followed by Tukey’s test. (* = p < 0.05 and ** = p < 0.01. Error bars represent ± SEM)

**Figure 6.**
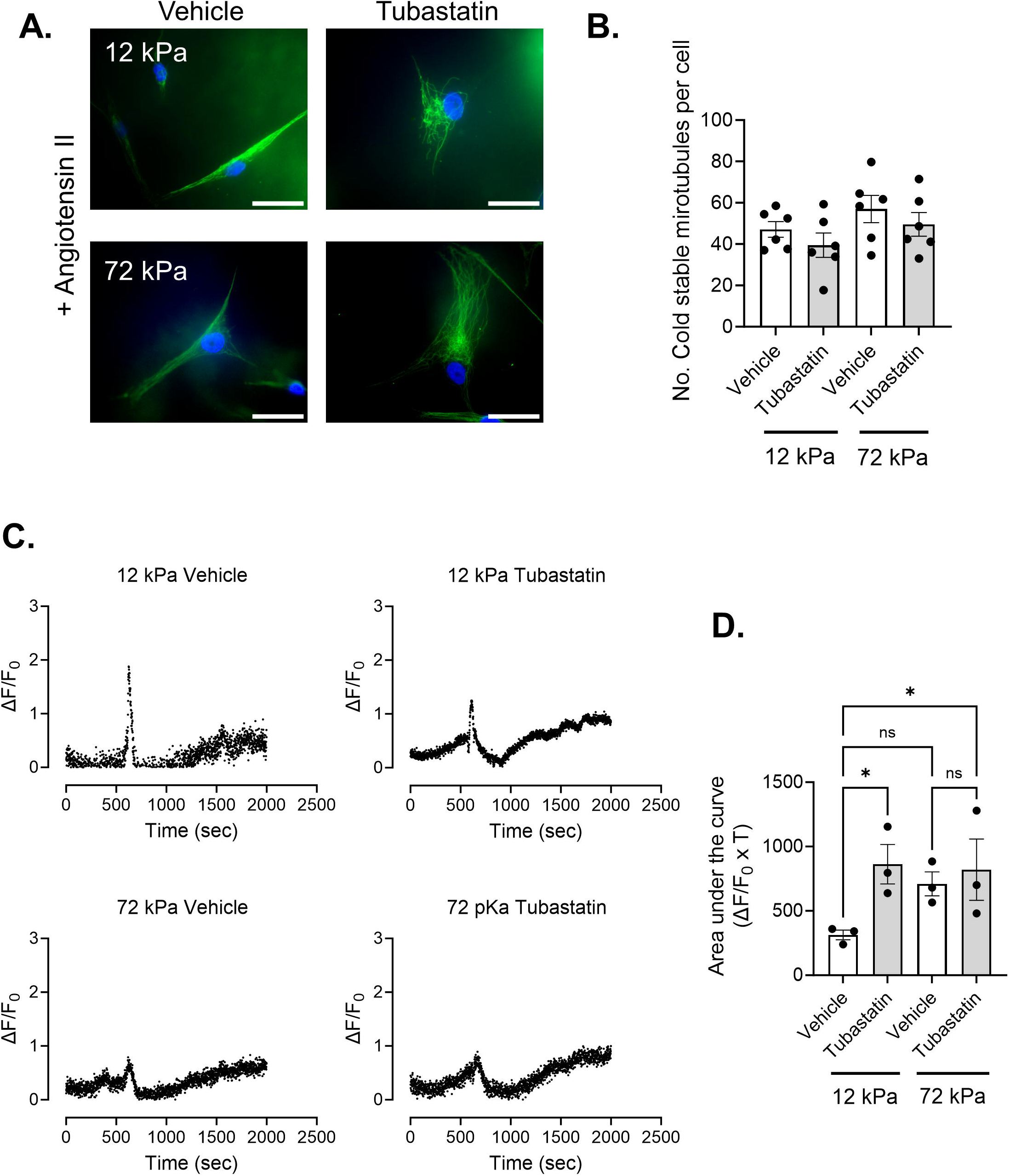
Tubastatin treatment alters Ca^2+^ flux in VSMCs. A) Representative images of isolated angiotensin II stimulated VSMCs pre-treated with vehicle control or tubastatin on 12 or 72 kPa polyacrylamide. Cold-stable microtubules, (α-tubulin, green) and nuclei (DAPI, blue). Scale bar = 50 μm. B) Graph shows number of cold-stable microtubules per cell on pliable and rigid hydrogels. Graph represents combined data from 6-independent experiments, analysing >60 cells per condition, and black dots mark mean data for each independent repeat. C) Graphs show representative ΔF/F_0_ values over time for vehicle control and tubastatin pre-treated angiotensin II stimulated Fluo-4 loaded VMSCs on 12 and 72 kPa hydrogels. Each graph represents the mean combined data of 9 individual fields of view combined from 3 individual experiments. D) Graph shows area under the curve values (ΔF/F_0_ x T). Mean data for individual repeats of the 3 independent experiments are shown as black dots. Significance was determined using two-way ANOVA followed by Tukey’s multiple comparison test. (* = p < 0.05. Error bars represent ± SEM)

## Discussion

Despite much research, our understanding of the mechanisms driving VSMC dysfunction and its contribution to decreased aortic compliance in ageing and CV disease remains limited. Previous studies have shown that enhanced matrix stiffness promotes the dedifferentiation of VSMCs, downregulating contractile markers whilst increasing the expression of proliferative genes (Brown et al., 2010; Nagayama and Nishimiya, 2020; Sazonova et al., 2015; Xie et al., 2018). Increases VSMC migrational capacity, adhesion, proliferation and volume have also been reported (Brown et al., 2010; Krajnik et al., 2023; Nagayama and Nishimiya, 2020; Rickel et al., 2020; Sazonova et al., 2015; Wong et al., 2003). Furthermore, in response to matrix stiffness, VSMC reorganise their actin cytoskeleton and generate enhanced traction stresses, a finding we recapitulate in this study(Brown et al., 2010; Petit et al., 2019; Sanyour et al., 2019; Sazonova et al., 2015).

Our findings show that changes in the tensegrity equilibrium defines VSMC behaviour in response to matrix rigidity. In this model, the microtubules serve as compression bearing struts that resist actomyosin generated strain (Brangwynne et al., 2006; Stamenović, 2005). Healthy VSMC behaviour is therefore a balance between microtubule stability and actomyosin activity. Actomyosin generated tension drives VSMC contraction and microtubules maintain VSMC morphology, protecting against strain induced cellular damage (Stamenović, 2005). The tensegrity model predicts that disruption of the microtubule cytoskeleton alters the equilibrium of these stresses and results in increased VSMC traction stress generation, a hypothesis previously confirmed by both wire myography and traction force microscopy (Ahmed et al., 2021; Zhang et al., 2000). We show that in rigid environments, the VSMC tensegrity equilibrium becomes unbalanced. Contractile agonist stimulation increases VSMC traction stress generation and reduces the number of cold-stable microtubules, resulting in deregulating VSMC morphology and enlargement of VSMC volume. Importantly, the altered tensegrity equilibrium can be induced by microtubule destabilising agents on pliable hydrogels and restored by treatment with agents that stabilise microtubules on rigid hydrogels. These data suggest that microtubules and the tensegrity equilibrium are key regulators of VSMC function and dysfunction.

Given the prevalence of increased aortic wall stiffness in ageing and disease, our findings suggest that altered tensegrity equilibrium is a key driver of VSMC dysfunction under these conditions (Tsamis et al., 2013). We propose that changing this equilibrium could be a mechanistic target for manipulating VSMC function. Using microtubule agents clinically is not a viable option to modulate this equilibrium, so a better understanding of the molecular mechanisms that regulate this equilibrium is now required to test this idea further.

In this study, we used Ca^2+^ flux as a readout of stretch-activated channel function. One of these, piezo1, is emerging as an important regulator of VSMC dysfunction (Johnson et al., 2024; Luu et al., 2023; Qian et al., 2022; Retailleau et al., 2015; Swiatlowska et al., 2023). Our findings further implicate piezo1, as a critical regulator of VSMC behaviour and suggest that changes in piezo1 opening drive altered VSMC behaviour in response to enhanced matrix stiffness. In this study, we show that VSMCs on rigid hydrogels display prolonged Ca^2+^ flux, increased traction stresses and reduced microtubule stability. The relationship between Ca^2+^ flux and microtubule stability were reciprocal, and changes in microtubule stability altered Ca^2+^ flux. We have previously shown that changes in microtubule stability also altered VSMC traction stress generation. Although untested, we predict that microtubule stability alters the ability of VSMCs to withstand the deformational forces placed on the plasma membrane, resulting in changes in membrane tension that control stretch-activated ion channel opening. Therefore, stretch-activated ion channel mediated Ca^2+^ influx serves as a mechanistic switch that triggers loss of the microtubule bearing struts and shifts the VSMC tensegrity equilibrium. We still do not understand the precise mechanisms that drive piezo1 mediated microtubule destabilisation. Millimolar Ca^2+^ concentrations spontaneously induce microtubule destabilisation in vitro, but cellular concentrations are much lower than this. We envisage 3 possibilities: 1) increased Ca^2+^ flux induces changes in the ability of microtubule stabilising proteins to associate with microtubules, resulting in destabilisation; 2) increased mechanical damage induces microtubule catastrophe; or 3) increased Ca^2+^ reduces microtubule growth/polymerisation.

The role of acetylation in microtubule stability, rigidity and resistance to mechanical damage remains controversial with many conflicting reports. We show that increased microtubule K40 acetylation stimulated both enhanced Ca^2+^ flux and VSMC volume on pliable hydrogels. This suggests that changes in microtubule post translational modification is also a factor in regulating VSMC matrix rigidity response. Despite this finding, K40 acetylation remained unaltered in VSMCs on pliable and rigid hydrogels in our study. This suggests that K40 acetylation does not play a role in the altered microtubule stability observed during VSMC matrix rigidity response.

However, HDAC6 inhibition failed to alter microtubule stability and appeared to uncouple the tensegrity equilibrium from the Ca^2+^ flux response on pliable hydrogels. Despite this uncoupling, the increased Ca^2+^ appeared to activate downstream signalling to enhance VSMC volume. HDAC6 has numerous targets, and we predict that this uncoupling is being driven by increased acetylation of other targets that remain unknown. Importantly, these findings suggest that, in addition to the tensegrity equilibrium and our previously identified volume control pathway, other unidentified mechanisms also contribute to the VSMC response to matrix rigidity. Further research is now needed to identify these unknown mechanisms for a more complete understanding of the VSMC matrix rigidity response.

In the healthy aortic wall, VSMCs adopt a quiescent, contractile differentiated phenotype. During ageing and disease, VSMCs downregulate contractile marker expression and modulate their phenotype towards disease relevant phenotypes (Pedroza et al., 2020; Wirka et al., 2019; Worssam et al., 2023). An important limitation of our study is that we have used dedifferentiated isolated VSMCs. VSMCs are known to possess multiple stretch-activated ion channels, and these can vary depending on phenotype. We have previously shown that piezo1 is upregulated in disease relevant phenotypes (Johnson et al., 2024). It remains unknown whether the VSMC tensegrity equilibrium can be altered in healthy contractile differentiated VSMCs, but microtubule agents alter VSMC contractile function in isolated aortic rings, suggesting that this is the case (Zhang et al., 2000). In this study, we implicate piezo1 in the defective matrix rigidity induced tensegrity equilibrium of isolated VSMCs. However, we cannot rule out the possibility that other stretch ion channels can induce altered tensegrity equilibrium in other phenotypes. Despite this limitation, our findings support the notion that the VSMC tensegrity equilibrium is modifiable and can be targeted to restore VSMC morphology.

## Materials and Methods

### Polyacrylamide Hydrogel Preparation

Hydrogels were prepared as described previously (Johnson et al., 2024; Minaisah et al., 2016). Briefly, glass coverslips were activated by treating with (3-Aminopropyl)triethoxysilane for 2 minutes, washed 3x in dH_2_O, then fixed in 0.5% glutaraldehyde for 40 minutes. After fixation, coverslips were washed and left to air dry overnight. Polyacrylamide hydrogel buffer was comprised as follows: 12 kPa – 7.5 % acrylamide, 0.15 % bis-acrylamide in dH_2_O; 72 kPa – 10 % acrylamide, 0.5 % bis-acrylamide in dH_2_O. To prepare hydrogels for fabrication, the appropriate volume of buffer was supplemented with 10 % APS (1:100) and TEMED (1:1,000) then placed on a standard microscopy slide and covered by an activated coverslip. Once set, the hydrogels were washed 3x in dH_2_O, cross-linked with sulfo-SANPAH (1:3000) under UV illumination (365 nm) for 5 minutes, then functionalised with collagen I (0.1 mg/ml) for 10 minutes at room temperature. Hydrogel stiffness was previously confirmed using a JPK Nanowizrd-3 atomic force microscope(Porter et al., 2020).

### Vascular Smooth Muscle Cell Culture

Human adult aortic VSMCs (passages 3-10) were purchased from Cell Applications Inc. (354-05a). Standard VSMC culture was performed as previously described (Ahmed et al., 2021; Johnson et al., 2024). VSMCs were seeded onto polyacrylamide hydrogels in basal media (Cell Applications Inc, Cat# 310-500), 18 hours prior to the beginning of the experiment to induce quiescence. Briefly, VSMCs were pre-treated for 30 minutes, prior to co-treatment with the contractile agonist, angiotensin II (10 µM) for an additional 30 minutes. Experimental specific concentrations are provided in the corresponding figure legends. For experiments performed in growth media, VSMCs were seeded onto hydrogels and incubated overnight. Pharmacological agents were added and VSMCs were incubated for a further 18 hours prior to fixation. Please see Table 1 below for details of compounds used in this study.

**Table 1.** Concentrations and details of compounds used in this study.

| Compound | Product Code | Supplier | Vehicle | Concentration Range Tested | Working Concentration |
| --- | --- | --- | --- | --- | --- |
| Angiotensin II | A9525 | Merck | dH<sub>2</sub>O | | 10 μM |
| Colchicine | C9754 | Merck | DMSO | 0.1 - 1000nM | 100 nM |
| Paclitaxel | T7402 | Merck | DMSO | 0.001 - 10 nM | 1 nM |
| Nocodazole | ab120630 | Abcam | DMSO | 0.001 - 10 nM | - |
| Epothilone B | E2656 | Merck | DMSO | 0.001 - 10 μM | - |
| Tubastatin A | #6270 | Tocris | DMSO | 0.01 – 1 μM | 1 μM |
| A23187 | | Tocris | DMSO | | 20 μM |

### siRNA Knockdown

VSMC siRNA transfection was performed using HiPerFect (Qiagen) as per manufacturer’s instructions, the day before cells were seeded onto hydrogels. VSMCs were transfected with either scrambled siRNA control, piezo1, or HDAC6 targeting siRNA (listed below) oligonucleotides. The next afternoon, cells were seeded onto hydrogels as above, serum was withdrawn overnight to induce quiescence and the next morning VSMCs were stimulated with angiotensin II (10 µM) for 30 minutes prior to fixation and downstream immunofluorescent analysis.

### Piezo1 siRNA

siRNA #5 – CCGCGTCTTCCTTAGCCATTA

siRNA #7 – CGGCCGCCTCGTGGTCTACAA

### HDAC6 siRNA

siRNA #5 – CACCGTCAACGTGGCATGGAA

siRNA #10 - CCGGAGGGTCCTTATCGTAGA

### Western Blotting

Western blotting was performed as previously described (Ragnauth et al., 2010). When looking for piezo1 expression specifically, lysates were run on a TruPAGE precast 4-20 % gradient gel (Sigma-Aldrich) at 120 V for 2 hours. Protein was transferred onto PVDF membrane at 30 V for 3 hours prior to the membrane being blocked in 5 % milk/TBST. The following antibodies were used: anti-piezo1 (1:500) (Novus Cat# NBP1-78537, RRID:AB_11003149), anti-GAPDH (1:4000) (Cell Signaling Technology Cat# 2118, RRID:AB_561053), anti-acetyl K40 alpha tubulin (Cell signaling technology Cat #3971, RRID:AB_2210204), anti-total alpha tubulin (Cell Signaling Technology Cat#3873, RRID:AB_1904178), anti-HDAC6 (D2E5) (Cell Signaling Technology Cat# 7558, RRID:AB_10891804), anti-Rabbit-HRP (1:2000) (Sigma-Aldrich Cat# GENA934, RRID:AB_2722659) and anti-Mouse-HRP (1:2000) (Sigma-Aldrich Cat# AP160P, RRID:AB_92531). 12 v 72 kPa lysates were repeated in triplicate for each individual experimental repeat.

### Immunofluorescence and VSMC Area/Volume Analysis

Cells were fixed in 4% paraformaldehyde for 10 minutes, permeabilised with 0.5 % v/v NP40 for 5 minutes, then blocked with 3 % w/v BSA/PBS for 1 hour. Primary staining against lamin A/C (1:200) (Sigma-Aldrich Cat# SAB4200236, RRID:AB_10743057) was performed overnight at 4 ^D^C in 3% BSA/PBS. Secondary antibody staining was performed using the appropriate Alexa Fluor™ 488 antibody (1:400) (Thermo Fisher Scientific Cat# A-11001, RRID:AB_2534069) in the dark for 2 hours. F-actin was visualised using Rhodamine Phalloidin (1:400) (Thermo Fisher Scientific Cat# R145). Images were captured at 20x magnification using a Zeiss LSM980-Airyscan confocal microscope. Cell area and volume was measured using FIJI, open-source software as described previously (Johnson et al., 2024; Schindelin et al., 2012).

### Traction Force Microscopy

VSMCs were seeded onto polyacrylamide hydrogels containing 0.5 µm red fluorescent (580/605) FluoSpheres (1:1000) (Invitrogen). Following angiotensin II stimulation (30 minutes), cell lysis was achieved by the addition of 0.5 % v/v Triton X-100. Images were captured at 20x magnification before and after lysis at 2-minute intervals using a Zeiss Axio Observer live cell imaging system. Drift was corrected using the ImageJ StackReg plugin and traction force was calculated using an ImageJ plugin that measured FluoSphere displacement (Tseng et al., 2012). Briefly, bead displacement was measured using the first and last image of the movie sequence. The cell region was determined by overlaying the traction map with the phase image, selecting the cell traction region with an ROI and extracting the traction forces in each pixel using the XY coordinate function in FIJI (Ahmed et al., 2021; Porter et al., 2020).

### Cold-Stable Microtubule Stability Assay

Cold-stable microtubules were identified as per previous studies (Atkinson et al., 2018). Following treatment, cells were placed on ice for 15 minutes before being washed once with PBS and twice with PEM buffer (80 μM PIPES pH 6.8, 1 mM, EGTA, 1 mM, MgCl_2_, 0.5 %, Triton X-100, and 25 % (w/v) glycerol) for 3 minutes. Cells were fixed in ice-cold methanol for 20 minutes then blocked with 3 % BSA/PBS for 1 hour. Microtubules were visualised by staining for α-tubulin (1:200) (Cell Signalling Technology Cat# 3873, RRID:AB_1904178) whilst cell nuclei were visualised using a mounting medium containing DAPI. Images were captured at 40x magnification using a Zeiss AxioPlan 2ie microscope.

### Cell Viability Assay

Cell viability was determined using a RealTime-Glo™ MT Cell Viability Assay, as per manufactures instructions. Briefly, 5,000 cells per well seeded in a 96-well plate and exposed to a range of drug concentrations for 1 hour. Luminescence was subsequently measured using a Wallac EnVision 2103 Multilabel Reader (PerkinElmer).

### Fluo-4 Calcium imaging

Cells seeded on 12 and 72kPa hydrogels and incubated in basal medium for 48 hours. Cells were loaded with 3 µM Fluo-4 AM (ThermoFisher Scientific cat# F14201) diluted in basal media for 30 minutes. Paclitaxel, colchicine and tubastatin were used at working concentrations listed in Supplementary Table 1. Cells were co-treated with vehicle control or compounds of interest during the Fluo-4 loading step. Cells were washed in PBS and returned to basal media prior to imaging. Cells were placed onto a Zeiss Axiovert 200M inverted microscope stage and images were captured every 500 msec. Cells were imaged for 5 minutes prior to angiotensin II (10 µM) stimulation. Cells were subsequently imaged for a further 20 minutes after addition of angiotensin II. Finally, the ionophore A23187 was added for a further 5 minutes to confirm the calcium imaging had worked. Images were analysed in ImageJ by selecting a ROI and extracting the fluorescence intensities for each time point using the plot Z-axis profile option. Background was subtracted and the ΔF and ΔF/F_0_ values were calculated and analysed using GraphPad prism.

### Statistical Analysis

Statistical analysis was performed using GraphPad Prism 9. Results are presented as mean ± SEM, with individual data points shown. The number of independent repeats performed, and total number of cells analysed per experiment are detailed in the corresponding figure legend. To compare more than two conditions a one-way ANOVA was performed, with a Tukey’s multiple comparison post-hoc test performed. Concentration response curves are presented as mean ± SEM plotted on a semi-logarithmic scale. Comparisons between concentration ranges on different hydrogel stiffness were performed using a two-way ANOVA followed by Sidak’s post-hoc test. Differences between conditions were considered statistically significant when P < 0.05.

## Author Contributions

RTJ, FW, RS, TM, CT, AB, OS, YB and DTW were responsible for performing experiments and analysing data. DTW was responsible for the conceptualisation; RTJ, SB, YB and DTW were responsible for the methodology of the study. SB, CJM and YB reviewed and edited this manuscript. RTJ, SB, YB and DTW and YB were responsible for the experimental design, writing and editing of this manuscript. RTJ assisted with the writing the initial version of the manuscript.

## Acknowledgements

This work was funded by a Biotechnology and Biological Sciences Research Council Research Grant (BB/T007699/1) awarded to DTW and a UKRI Biotechnology and Biological Sciences Research Council Norwich Research Park Doctoral Training Partnership PhD Studentship awarded to FW (BB/T008717/1). RS was funded by a University of East Anglia Science Faculty PhD Studentship.

## Data Availability

The data that support the findings of this study are available from the corresponding author upon reasonable request.

## References

Afewerki, T., Ahmed, S., Warren, D., 2019. Emerging regulators of vascular smooth muscle cell migration. J Muscle Res Cell Motil 40, 185–196. 10.1007/s10974-019-09531-z

Ahmed, S., Johnson, R.T., Solanki, R., Afewerki, T., Wostear, F., Warren, D.T., 2021. Using polyacrylamide hydrogels to model physiological aortic stiffness reveals that microtubules are critical regulators of isolated smooth muscle cell morphology and contractility. 10.1101/2021.12.14.472278

Ahmed, S., Warren, D.T., 2018. Vascular smooth muscle cell contractile function and mechanotransduction. Vessel Plus 2, 36. 10.20517/2574-1209.2018.51

Atkinson, S.J., Gontarczyk, A.M., Alghamdi, A.A., Ellison, T.S., Johnson, R.T., Fowler, W.J., Kirkup, B.M., Silva, B.C., Harry, B.E., Schneider, J.G., Weilbaecher, K.N., Mogensen, M.M., Bass, M.D., Parsons, M., Edwards, D.R., Robinson, S.D., 2018. The β3-integrin endothelial adhesome regulates microtubule-dependent cell migration. EMBO Rep 19, e44578. 10.15252/embr.201744578

Brangwynne, C.P., MacKintosh, F.C., Kumar, S., Geisse, N.A., Talbot, J., Mahadevan, L., Parker, K.K., Ingber, D.E., Weitz, D.A., 2006. Microtubules can bear enhanced compressive loads in living cells because of lateral reinforcement. Journal of Cell Biology 173, 733–741. 10.1083/jcb.200601060

Brown, X.Q., Bartolak-Suki, E., Williams, C., Walker, M.L., Weaver, V.M., Wong, J.Y., 2010. Effect of substrate stiffness and PDGF on the behavior of vascular smooth muscle cells: implications for atherosclerosis. J Cell Physiol 225, 115–122. 10.1002/jcp.22202

Cao, G., Xuan, X., Hu, J., Zhang, R., Jin, H., Dong, H., 2022. How vascular smooth muscle cell phenotype switching contributes to vascular disease. Cell Communication and Signaling 20, 180. 10.1186/s12964-022-00993-2

Coleman, A.K., Joca, H.C., Shi, G., Lederer, W.J., Ward, C.W., 2021. Tubulin acetylation increases cytoskeletal stiffness to regulate mechanotransduction in striated muscle. Journal of General Physiology 153, e202012743. 10.1085/jgp.202012743

Eshun-Wilson, L., Zhang, R., Portran, D., Nachury, M.V., Toso, D.B., Löhr, T., Vendruscolo, M., Bonomi, M., Fraser, J.S., Nogales, E., 2019. Effects of α-tubulin acetylation on microtubule structure and stability. Proc. Natl. Acad. Sci. U.S.A. 116, 10366–10371. 10.1073/pnas.1900441116

Glasser, S.P., Arnett, D.K., McVeigh, G.E., Finkelstein, S.M., Bank, A.J., Morgan, D.J., Cohn, J.N., 1997. Vascular Compliance and Cardiovascular Disease: A Risk Factor or a Marker? American Journal of Hypertension 10, 1175–1189. 10.1016/S0895-7061(97)00311-7

Hayashi, K., Naiki, T., 2009. Adaptation and remodeling of vascular wall; biomechanical response to hypertension. Journal of the Mechanical Behavior of Biomedical Materials 2, 3–19. 10.1016/j.jmbbm.2008.05.002

Janke, C., Montagnac, G., 2017. Causes and Consequences of Microtubule Acetylation. Current Biology 27, R1287–R1292. 10.1016/j.cub.2017.10.044

Johnson, R.T., Solanki, R., Warren, D.T., 2021. Mechanical programming of arterial smooth muscle cells in health and ageing. Biophys Rev 13, 757–768. 10.1007/s12551-021-00833-6

Johnson, R.T., Solanki, R., Wostear, F., Ahmed, S., Taylor, J.C.K., Rees, J., Abel, G., McColl, J., Jørgensen, H.F., Morris, C.J., Bidula, S., Warren, D.T., 2024. Piezo1-mediated regulation of smooth muscle cell volume in response to enhanced extracellular matrix rigidity. British J Pharmacology bph.16294. 10.1111/bph.16294

Krajnik, A., Nimmer, E., Brazzo, J.A., Biber, J.C., Drewes, R., Tumenbayar, B.-I., Sullivan, A., Pham, K., Krug, A., Heo, Y., Kolega, J., Heo, S.-J., Lee, K., Weil, B.R., Kim, D.-H., Gupte, S.A., Bae, Y., 2023. Survivin regulates intracellular stiffness and extracellular matrix production in vascular smooth muscle cells. APL Bioengineering 7, 046104. 10.1063/5.0157549

Lacolley, P., Regnault, V., Laurent, S., 2020. Mechanisms of Arterial Stiffening. Arteriosclerosis, Thrombosis, and Vascular Biology 40, 1055–1062. 10.1161/ATVBAHA.119.313129

Luu, N., Bajpai, A., Li, R., Park, S., Noor, M., Ma, X., Chen, W., 2023. Aging-associated decline in vascular smooth muscle cell mechanosensation is mediated by Piezo1 channel. Aging Cell e14036. 10.1111/acel.14036

Michaels, T.C., Feng, S., Liang, H., Mahadevan, L., 2020. Mechanics and kinetics of dynamic instability. eLife 9, e54077. 10.7554/eLife.54077

Minaisah, R.-M., Cox, S., Warren, D.T., 2016. The Use of Polyacrylamide Hydrogels to Study the Effects of Matrix Stiffness on Nuclear Envelope Properties, in: Shackleton, S., Collas, P., Schirmer, E.C. (Eds.), The Nuclear Envelope, Methods in Molecular Biology. Springer New York, New York, NY, pp. 233–239. 10.1007/978-1-4939-3530-7_15

Mitchell, G.F., Hwang, S.-J., Vasan, R.S., Larson, M.G., Pencina, M.J., Hamburg, N.M., Vita, J.A., Levy, D., Benjamin, E.J., 2010. Arterial Stiffness and Cardiovascular Events. Circulation 121, 505– 511. 10.1161/CIRCULATIONAHA.109.886655

Nagayama, K., Nishimiya, K., 2020. Moderate substrate stiffness induces vascular smooth muscle cell differentiation through cellular morphological and tensional changes. Biomed Mater Eng 31, 157–167. 10.3233/BME-201087

Osseni, A., Ravel-Chapuis, A., Thomas, J.-L., Gache, V., Schaeffer, L., Jasmin, B.J., 2020. HDAC6 regulates microtubule stability and clustering of AChRs at neuromuscular junctions. Journal of Cell Biology 219, e201901099. 10.1083/jcb.201901099

Owens, G.K., Schwartz, S.M., 1983. Vascular smooth muscle cell hypertrophy and hyperploidy in the Goldblatt hypertensive rat. Circulation Research 53, 491–501. 10.1161/01.RES.53.4.491

Pedroza, A.J., Tashima, Y., Shad, R., Cheng, P., Wirka, R., Churovich, S., Nakamura, K., Yokoyama, N., Cui, J.Z., Iosef, C., Hiesinger, W., Quertermous, T., Fischbein, M.P., 2020. Single-Cell Transcriptomic Profiling of Vascular Smooth Muscle Cell Phenotype Modulation in Marfan Syndrome Aortic Aneurysm. Arteriosclerosis, Thrombosis, and Vascular Biology 40, 2195– 2211. 10.1161/ATVBAHA.120.314670

Petit, C., Guignandon, A., Avril, S., 2019. Traction Force Measurements of Human Aortic Smooth Muscle Cells Reveal a Motor-Clutch Behavior. Molecular and Cellular Biomechanics. 10.32604/mcb.2019.06415

Porter, L., Minaisah, R.-M., Ahmed, S., Ali, S., Norton, R., Zhang, Q., Ferraro, E., Molenaar, C., Holt, M., Cox, S., Fountain, S., Shanahan, C., Warren, D., 2020. SUN1/2 Are Essential for RhoA/ROCK-Regulated Actomyosin Activity in Isolated Vascular Smooth Muscle Cells. Cells 9, 132. 10.3390/cells9010132

Qian, W., Hadi, T., Silvestro, M., Ma, X., Rivera, C.F., Bajpai, A., Li, R., Zhang, Z., Qu, H., Tellaoui, R.S., Corsica, A., Zias, A.L., Garg, K., Maldonado, T., Ramkhelawon, B., Chen, W., 2022. Microskeletal stiffness promotes aortic aneurysm by sustaining pathological vascular smooth muscle cell mechanosensation via Piezo1. Nat Commun 13, 512. 10.1038/s41467-021-27874-5

Ragnauth, C.D., Warren, D.T., Liu, Y., McNair, R., Tajsic, T., Figg, N., Shroff, R., Skepper, J., Shanahan, C.M., 2010. Prelamin A Acts to Accelerate Smooth Muscle Cell Senescence and Is a Novel Biomarker of Human Vascular Aging. Circulation 121, 2200–2210. 10.1161/CIRCULATIONAHA.109.902056

Retailleau, K., Duprat, F., Arhatte, M., Ranade, S.S., Peyronnet, R., Martins, J.R., Jodar, M., Moro, C., Offermanns, S., Feng, Y., Demolombe, S., Patel, A., Honoré, E., 2015. Piezo1 in Smooth Muscle Cells Is Involved in Hypertension-Dependent Arterial Remodeling. Cell Reports 13, 1161–1171. 10.1016/j.celrep.2015.09.072

Rickel, A.P., Sanyour, H.J., Leyda, N.A., Hong, Z., 2020. Extracellular Matrix Proteins and Substrate Stiffness Synergistically Regulate Vascular Smooth Muscle Cell Migration and Cortical Cytoskeleton Organization. ACS Appl. Bio Mater. 3, 2360–2369. 10.1021/acsabm.0c00100

Rizzoni, D., Porteri, E., Guefi, D., Piccoli, A., Castellano, M., Pasini, G., Muiesan, M.L., Mulvany, M.J., Rosei, E.A., 2000. Cellular Hypertrophy in Subcutaneous Small Arteries of Patients With Renovascular Hypertension. Hypertension 35, 931–935. 10.1161/01.HYP.35.4.931

Sanders, K.M., 2001. Invited Review: Mechanisms of calcium handling in smooth muscles. Journal of Applied Physiology 91, 1438–1449. 10.1152/jappl.2001.91.3.1438

Sanyour, H.J., Li, N., Rickel, A.P., Childs, J.D., Kinser, C.N., Hong, Z., 2019. Membrane cholesterol and substrate stiffness co-ordinate to induce the remodelling of the cytoskeleton and the alteration in the biomechanics of vascular smooth muscle cells. Cardiovasc Res 115, 1369– 1380. 10.1093/cvr/cvy276

Sazonova, O.V., Isenberg, B.C., Herrmann, J., Lee, K.L., Purwada, A., Valentine, A.D., Buczek-Thomas, J.A., Wong, J.Y., Nugent, M.A., 2015. Extracellular matrix presentation modulates vascular smooth muscle cell mechanotransduction. Matrix Biology 41, 36–43. 10.1016/j.matbio.2014.11.001

Schiffrin, E.L., 2012. Vascular Remodeling in Hypertension. Hypertension 59, 367–374. 10.1161/HYPERTENSIONAHA.111.187021

Schindelin, J., Arganda-Carreras, I., Frise, E., Kaynig, V., Longair, M., Pietzsch, T., Preibisch, S., Rueden, C., Saalfeld, S., Schmid, B., Tinevez, J.-Y., White, D.J., Hartenstein, V., Eliceiri, K., Tomancak, P., Cardona, A., 2012. Fiji: an open-source platform for biological-image analysis. Nat Methods 9, 676–682. 10.1038/nmeth.2019

Sehgel, N.L., Vatner, S.F., Meininger, G.A., 2015. “Smooth Muscle Cell Stiffness Syndrome”— Revisiting the Structural Basis of Arterial Stiffness. Frontiers in Physiology 6.

Sehgel, N.L., Zhu, Y., Sun, Z., Trzeciakowski, J.P., Hong, Z., Hunter, W.C., Vatner, D.E., Meininger, G.A., Vatner, S.F., 2013. Increased vascular smooth muscle cell stiffness: a novel mechanism for aortic stiffness in hypertension. American Journal of Physiology-Heart and Circulatory Physiology 305, H1281–H1287. 10.1152/ajpheart.00232.2013

Shida, T., Cueva, J.G., Xu, Z., Goodman, M.B., Nachury, M.V., 2010. The major α-tubulin K40 acetyltransferase αTAT1 promotes rapid ciliogenesis and efficient mechanosensation. Proc. Natl. Acad. Sci. U.S.A. 107, 21517–21522. 10.1073/pnas.1013728107

Song, Y., Brady, S.T., 2015. Post-translational modifications of tubulin: pathways to functional diversity of microtubules. Trends in Cell Biology 25, 125–136. 10.1016/j.tcb.2014.10.004

Stamenović, D., 2005. Microtubules may harden or soften cells, depending of the extent of cell distension. Journal of Biomechanics 38, 1728–1732. 10.1016/j.jbiomech.2004.07.016

Steucke, K.E., Win, Z., Stemler, T.R., Walsh, E.E., Hall, J.L., Alford, P.W., 2017. Empirically Determined Vascular Smooth Muscle Cell Mechano-Adaptation Law. Journal of Biomechanical Engineering 139, 071005. 10.1115/1.4036454

Swiatlowska, P., Tipping, W., Marhuenda, E., Severi, P., Fomin, V., Yang, Z., Xiao, Q., Graham, D., Shanahan, C., Iskratsch, T., 2023. Hypertensive Pressure Mechanosensing Alone Triggers Lipid Droplet Accumulation and Transdifferentiation of Vascular Smooth Muscle Cells to Foam Cells. Advanced Science 2308686. 10.1002/advs.202308686

Tsamis, A., Krawiec, J.T., Vorp, D.A., 2013. Elastin and collagen fibre microstructure of the human aorta in ageing and disease: a review. Journal of The Royal Society Interface 10, 20121004. 10.1098/rsif.2012.1004

Wirka, R.C., Wagh, D., Paik, D.T., Pjanic, M., Nguyen, T., Miller, C.L., Kundu, R., Nagao, M., Coller, J., Koyano, T.K., Fong, R., Woo, Y.J., Liu, B., Montgomery, S.B., Wu, J.C., Zhu, K., Chang, R., Alamprese, M., Tallquist, M.D., Kim, J.B., Quertermous, T., 2019. Atheroprotective roles of smooth muscle cell phenotypic modulation and the TCF21 disease gene as revealed by single-cell analysis. Nat Med 25, 1280–1289. 10.1038/s41591-019-0512-5

Wong, J.Y., Velasco, A., Rajagopalan, P., Pham, Q., 2003. Directed Movement of Vascular Smooth Muscle Cells on Gradient-Compliant Hydrogels. Langmuir 19, 1908–1913. 10.1021/la026403p

Worssam, M.D., Lambert, J., Oc, S., Taylor, J.C.K., Taylor, A.L., Dobnikar, L., Chappell, J., Harman, J.L., Figg, N.L., Finigan, A., Foote, K., Uryga, A.K., Bennett, M.R., Spivakov, M., Jørgensen, H.F., 2023. Cellular mechanisms of oligoclonal vascular smooth muscle cell expansion in cardiovascular disease. Cardiovasc Res 119, 1279–1294. 10.1093/cvr/cvac138

Xie, S.-A., Zhang, T., Wang, J., Zhao, F., Zhang, Y.-P., Yao, W.-J., Hur, S.S., Yeh, Y.-T., Pang, W., Zheng, L.-S., Fan, Y.-B., Kong, W., Wang, X., Chiu, J.-J., Zhou, J., 2018. Matrix stiffness determines the phenotype of vascular smooth muscle cell in vitro and in vivo: Role of DNA methyltransferase 1. Biomaterials 155, 203–216. 10.1016/j.biomaterials.2017.11.033

Xu, Z., Schaedel, L., Portran, D., Aguilar, A., Gaillard, J., Marinkovich, M.P., Théry, M., Nachury, M.V., 2017. Microtubules acquire resistance from mechanical breakage through intralumenal acetylation. Science 356, 328–332. 10.1126/science.aai8764

Zhang, D., Jin, N., Rhoades, R.A., Yancey, K.W., Swartz, D.R., 2000. Influence of microtubules on vascular smooth muscle contraction. J Muscle Res Cell Motil 21, 293–300. 10.1023/A:1005600118157

Zhang, Y., Griendling, K.K., Dikalova, A., Owens, G.K., Taylor, W.R., 2005. Vascular Hypertrophy in Angiotensin II–Induced Hypertension Is Mediated by Vascular Smooth Muscle Cell–Derived H2O2. Hypertension 46, 732–737. 10.1161/01.HYP.0000182660.74266.6d

