## Supplementary Figures for "A microtubule stability switch alters isolated vascular smooth muscle calcium flux in response to matrix rigidity"

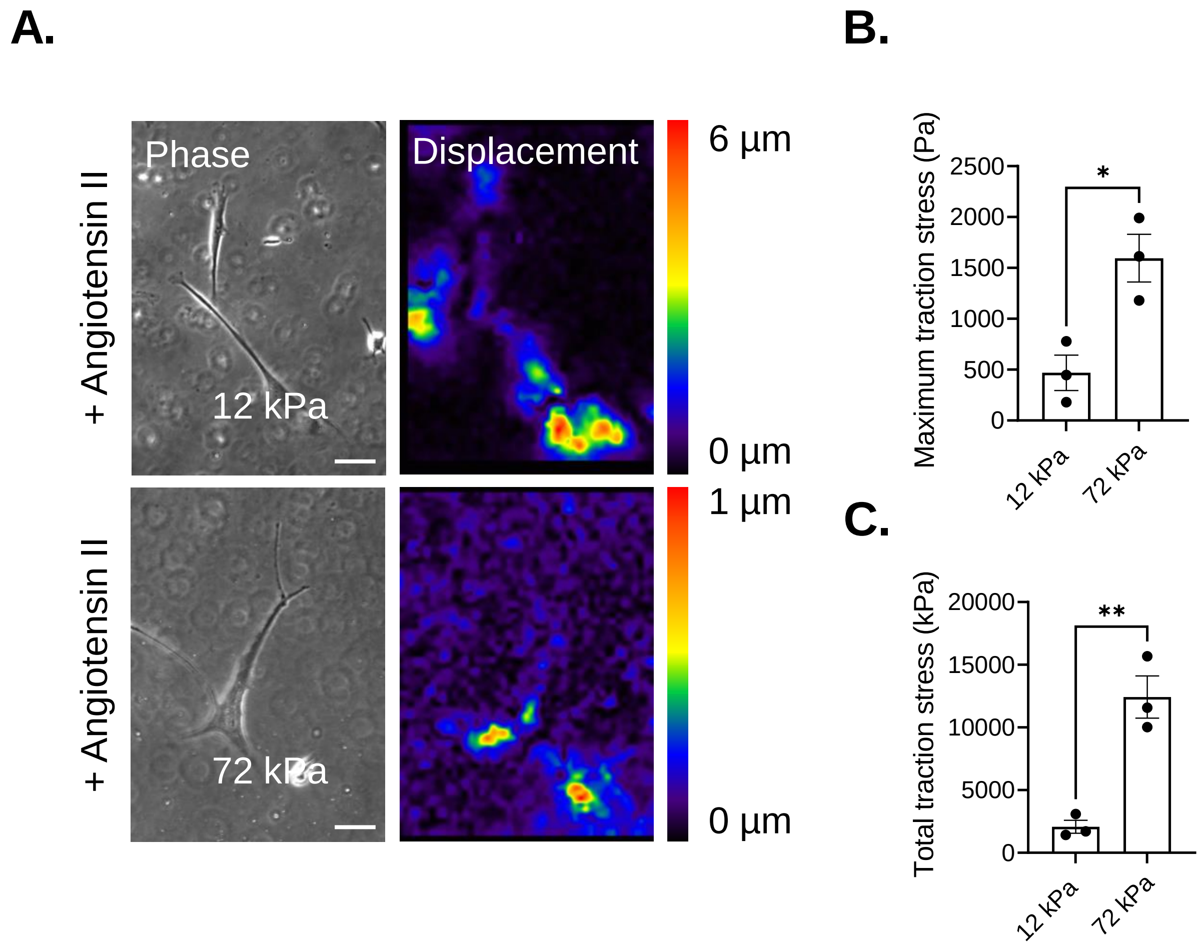


**Supplementary Figure S1. VSMCs generate enhanced traction stress on rigid hydrogels.** A) Representative phase images and bead displacement heat maps of AngII (10 µM) stimulated VSMCs cultured on 12 or 72 kPa polyacrylamide hydrogels. Scale bar = 100 µm. Graphs show B) maximum and C) total traction stress generation, representative of 3 independent experiments, with ≥38 cells analysed per condition. Black dots represent mean data for each individual experimental repeat. Significance determined using unpaired student t-Test. * = $p < 0.05$ and ** = $p < 0.01$. Error bars represent ± SEM.

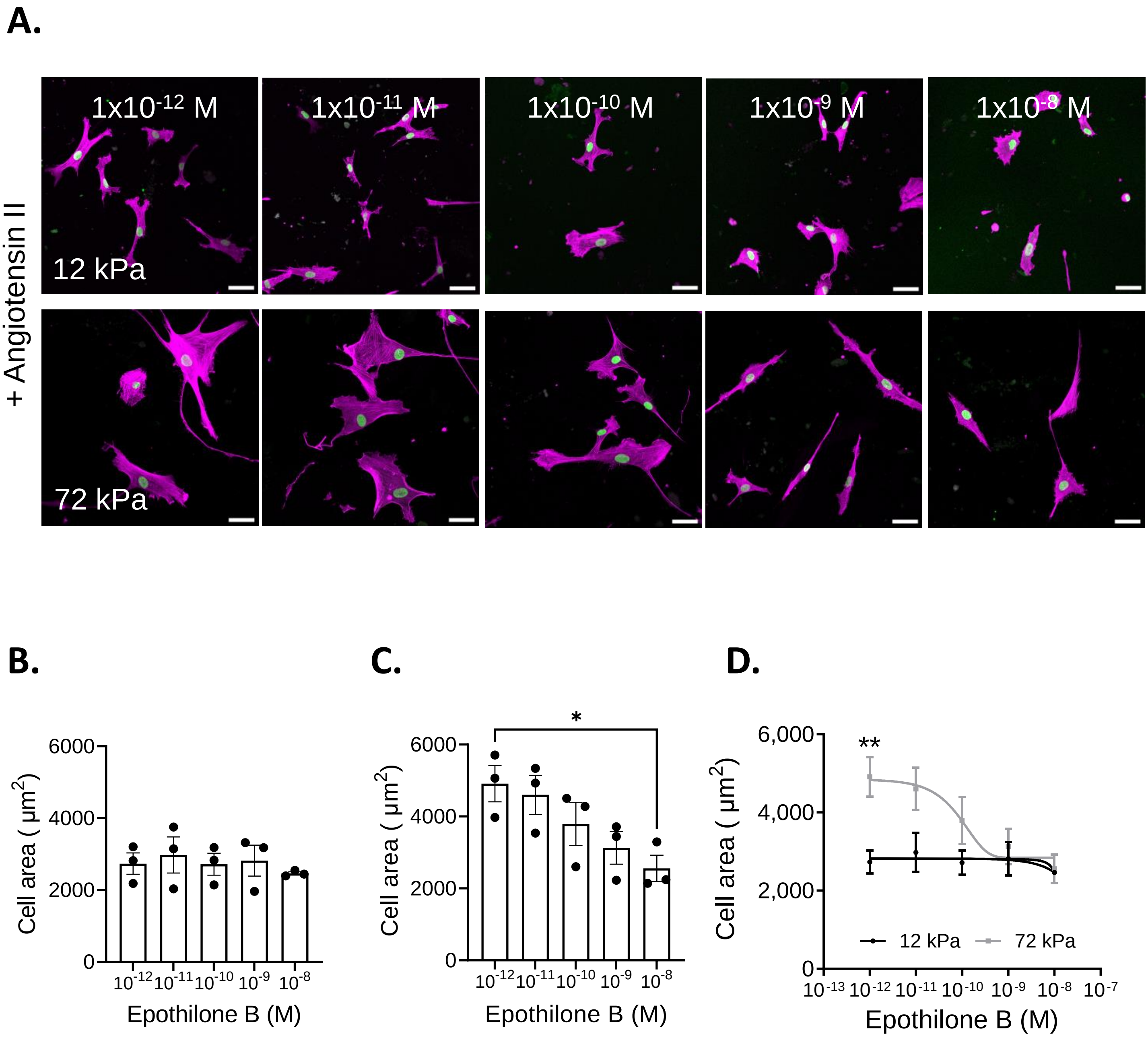


**Supplementary Figure S2. Epothilone treatment blocks increased VSMC area response on rigid hydrogels.** A) Representative images of VSMCs pre-treated with increasing concentrations of epothilone B on 12 kPa and 72 kPa hydrogels. Purple = Rhodamine Phalloidin, green = DAPI and Scale bar = 50 µm. Graphs show VSMC area on B) 12 kPa and C) 72 kPa hydrogels representative of 3 independent experiments with ≥53 cells analysed per condition. Black dots represent mean data from each individual experimental repeat. Significance determined using a one-way ANOVA followed by Tukey's test. D) Comparison of VSMC response to epothilone B pre-treatment on 12 and 72 kPa hydrogels. Data is expressed as the mean of the means calculated from 3 independent experiments; significance determined using a two-way ANOVA followed by Sidak's test. (* = $p < 0.05$ and ** = $p < 0.01$. Error bars represent ± SEM).

**Supplementary Figure S2**

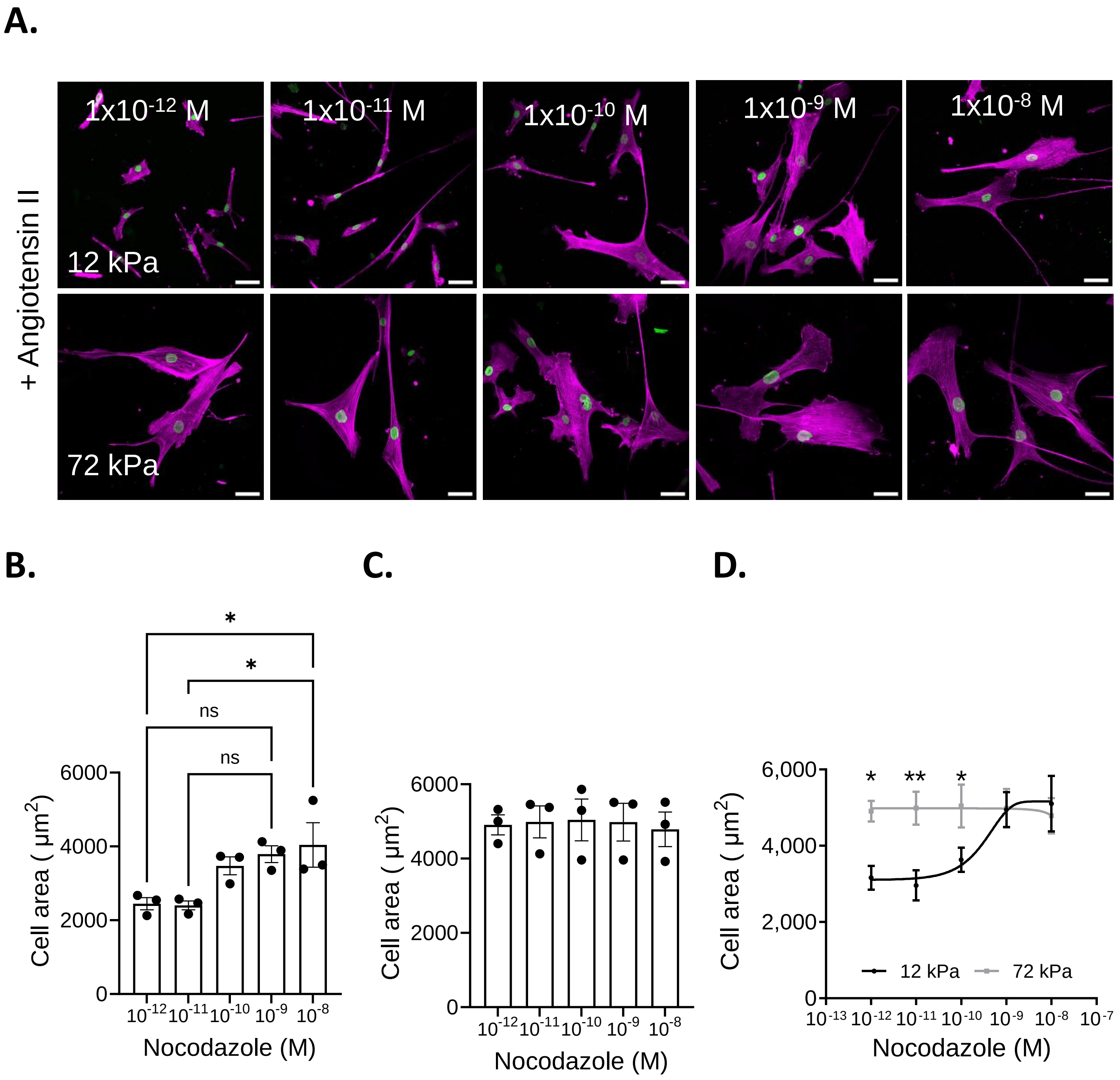


**Supplementary Figure S3. Nocodazole treatment increases VSMC area on pliable hydrogels.** A) Representative images of VSMCs pre-treated with increasing concentrations of nocodazole on 12 kPa and 72 kPa hydrogels. Purple = Rhodamine Phalloidin, green = DAPI and Scale bar = 50 µm. Graphs show VSMC area on B) 12 kPa and C) 72 kPa hydrogels representative of 3 independent experiments with ≥58 cells analysed per condition. Black dots represent mean data from each individual experimental repeat. Significance determined using a one-way ANOVA followed by Tukey's test. D) Comparison of VSMC response to nocodazole pre-treatment on 12 and 72 kPa hydrogels. Data is expressed as the mean of the means calculated from 3 independent experiments; significance determined using a two-way ANOVA followed by Sidak's test. (* = $p < 0.05$ and ** = $p < 0.01$. Error bars represent ± SEM).

**Supplementary Figure S3**

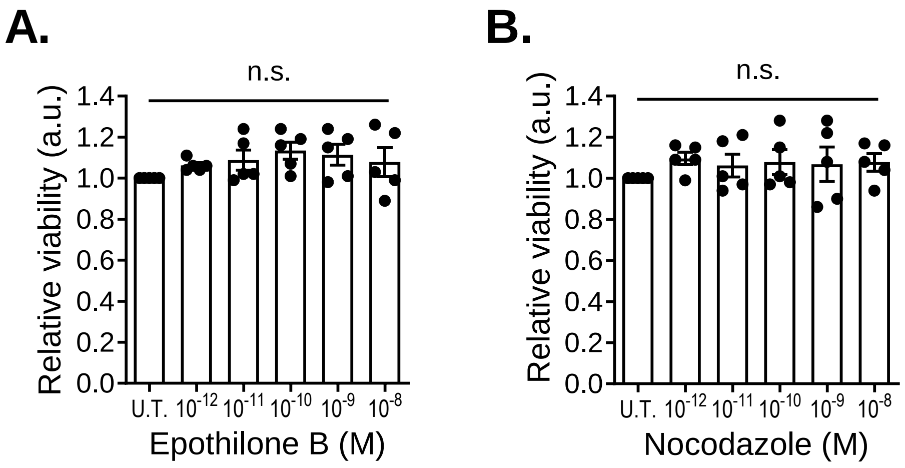


**Supplementary Figure S4. Regulators of microtubule stability have no effect on VSMC viability.** Relative VSMC viability following a 1-hour treatment with increasing concentrations of A) epothilone B and B) nocodazole. U.T. = Untreated. Data is representative of 5 independent experiments; significance determined using a one-way ANOVA followed by Tukey's test. (n.s. = *non-significant*, error bars represent ± SEM).

**Supplementary Figure S4**

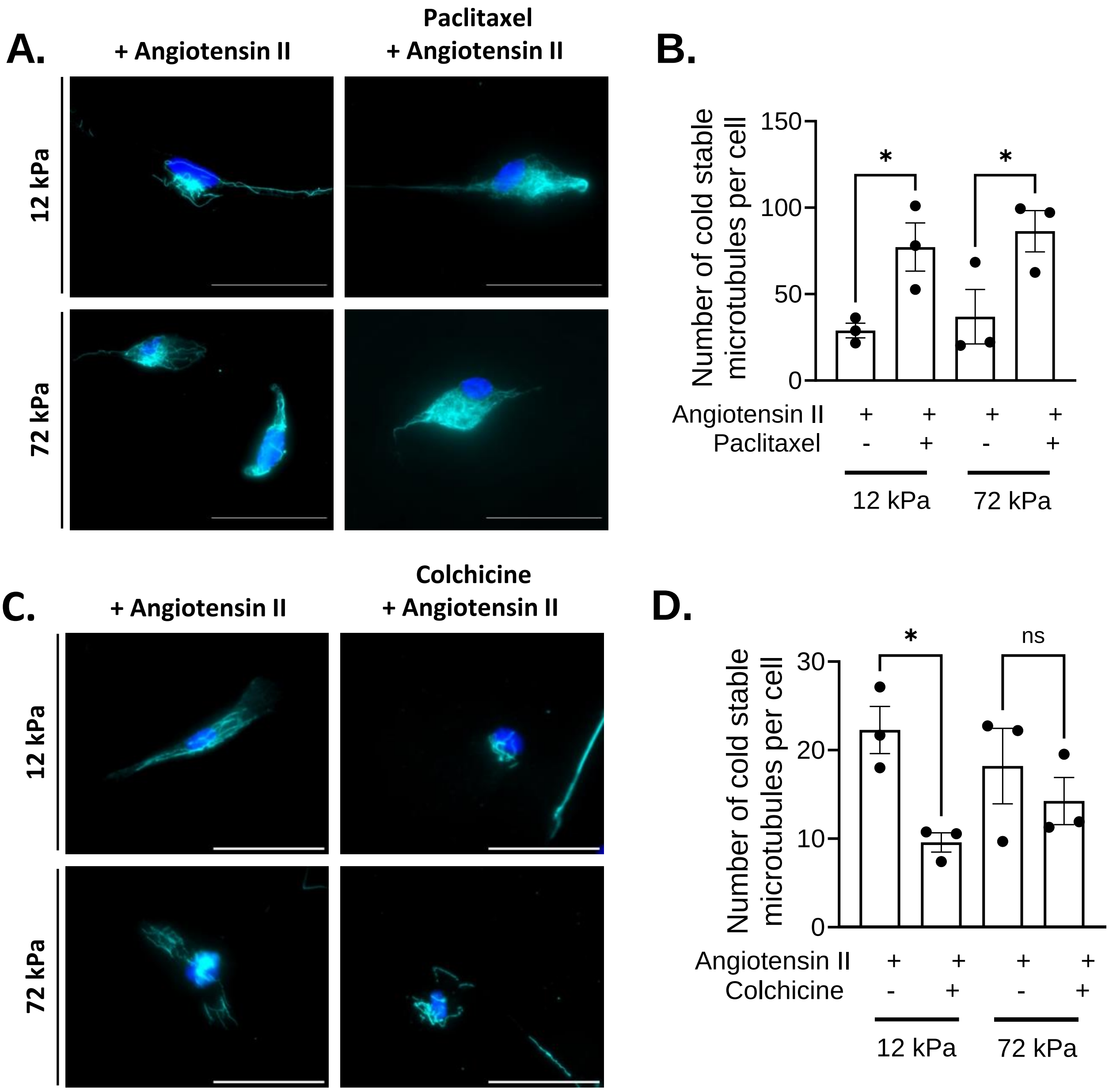


**Supplementary Figure S5. Microtubule targeting agents regulate microtubule stability in isolated VSMCs.** Representative images of isolated VSMCs cultured on 12 or 72 kPa polyacrylamide hydrogels pretreated with A) paclitaxel or C) colchicine prior to angiotensin II stimulation. Cold-stable microtubules, (α-tubulin, aqua) and nuclei (DAPI, blue). Scale bar = 50 µm. Graphs show number of cold-stable microtubules per cell after B) paclitaxel treatment, representative of 3 independent experiments, with ≥65 cells analysed per condition; and D) colchicine treatment, representative of 3 independent experiments, with ≥82 cells analysed per condition. Black dots represent mean data for each individual experimental repeat. Significance determined using a one-way ANOVA followed by Sidak's test. (* = $p < 0.05$, error bars represent ± SEM).

**Supplementary Figure S5**

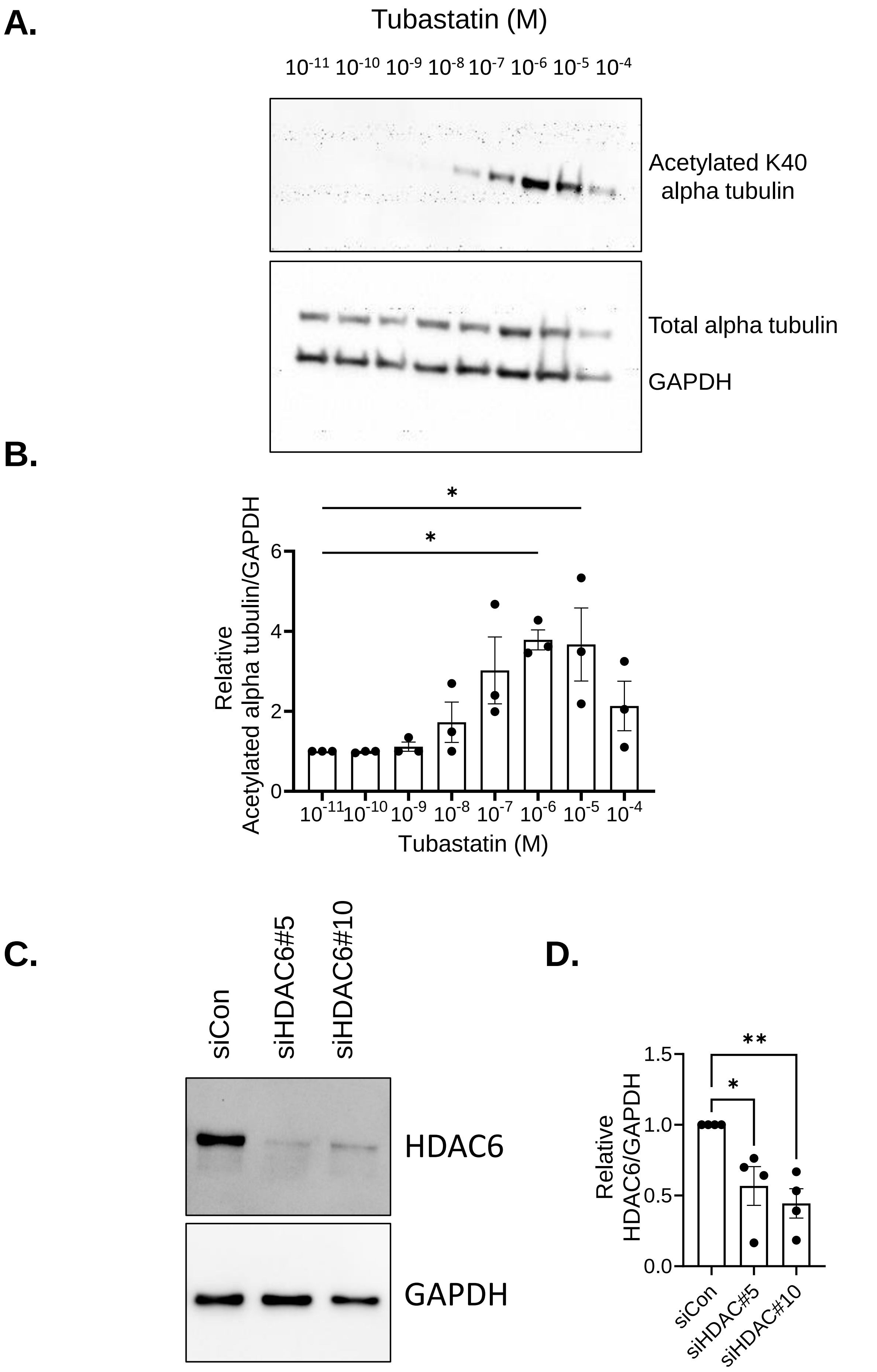


**Supplementary Figure S6. Conformation of tubastatin function and HDAC6 depletion in VSMCs. A)** Representative WB of acetyl-alpha tubulin levels in isolated VSMCs treated with a concentration range of tubastatin for 1 hour. **B)** Graph shows the relative acetyl-alpha tubulin/GAPDH level determine by densitometry and represents the combined data of 3-independent experiments. Black dots represent the mean data of each individual experimental repeat. Significance determined using a one-way ANOVA followed by Sidak's test. (* = $p < 0.05$, error bars represent ± SEM). C) Representative WB of HDAC6 levels in scrambled (siCon) and siHDAC6 specific (siHDAC6#5 and siHDAC6#10) siRNA. D) Graph shows the relative HDAC6/GAPDH level determined by densitometry and represents the combined data of 4-independent experiments. Black dots represent the mean data of each individual experimental repeat. Significance determined using a one-way ANOVA followed by Sidak's test. (* = $p < 0.05$, error bars represent ± SEM).

**Supplementary Figure S6**

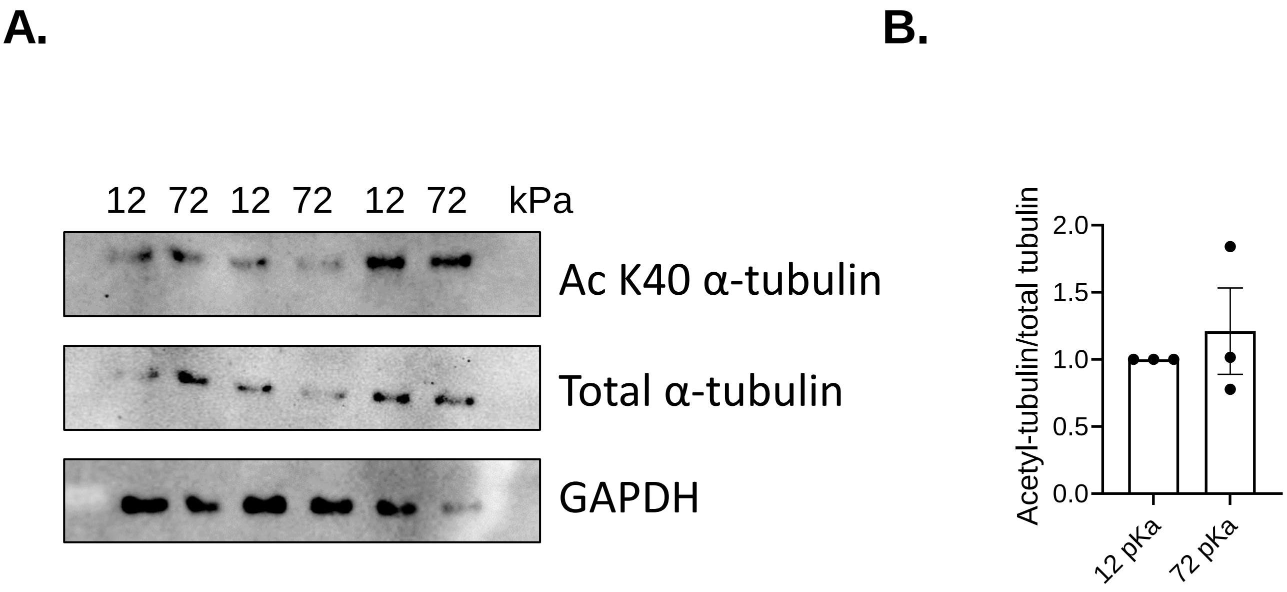


**Supplementary Figure S7. Matrix stiffness does not alter acetyl-alpha tubulin levels in VSMCs. A)** Representative WB of acetyl-alpha tubulin levels in isolated VSMCs grown on 12 and 72 kPa hydrogels for 3 days. **B)** Graph shows the relative acetyl-alpha tubulin/total tubulin level determine by densitometry and represents the combined data of 3-independent experiments. Black dots represent the mean data of each individual experimental repeat. Significance determined using a one-way ANOVA followed by Sidak's test. (* = $p < 0.05$, error bars represent ± SEM).

**Blot transparency**

**Figure 1C uncropped WBs.** Some lanes removed as loaded with lysates that are not part of this study. This extended image contains an untransfected control and an addition piezo1 targeting siRNA (siP#10)

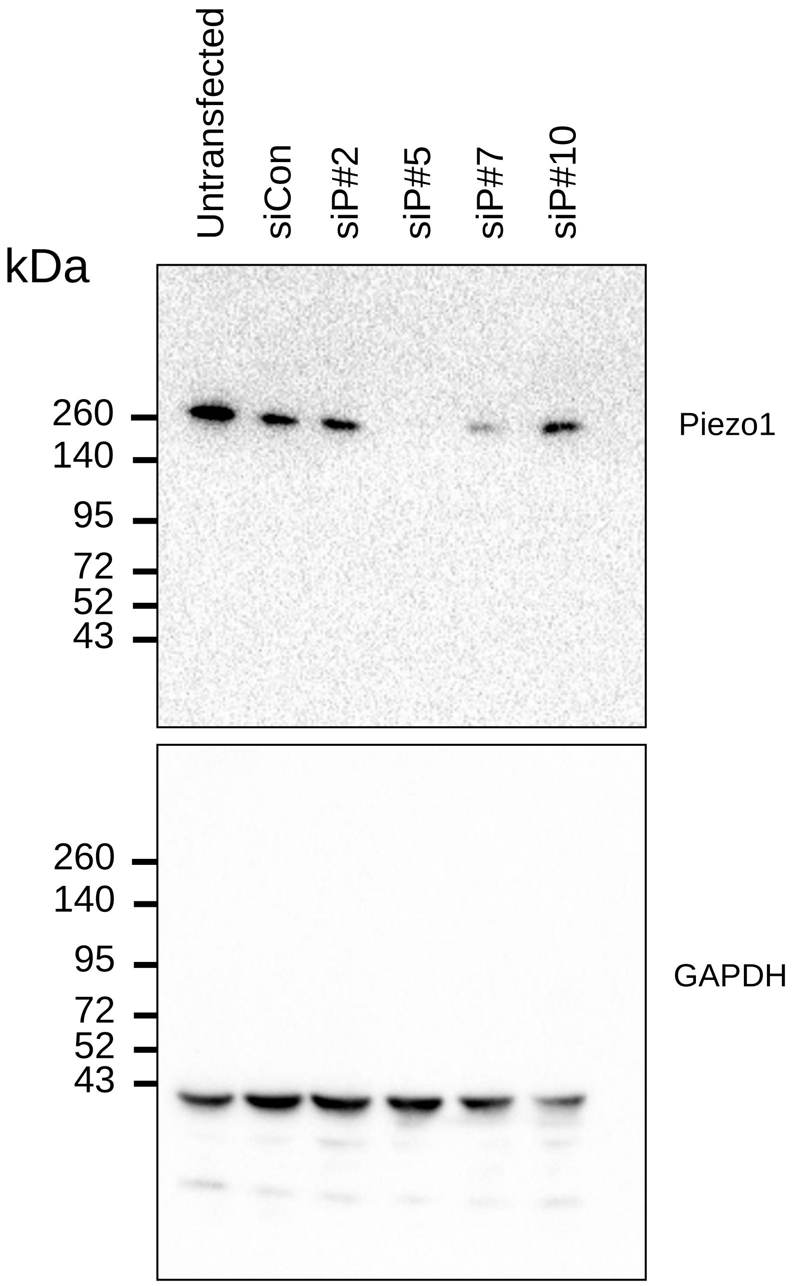

**Supplementary Figure S6A uncropped WBs.**

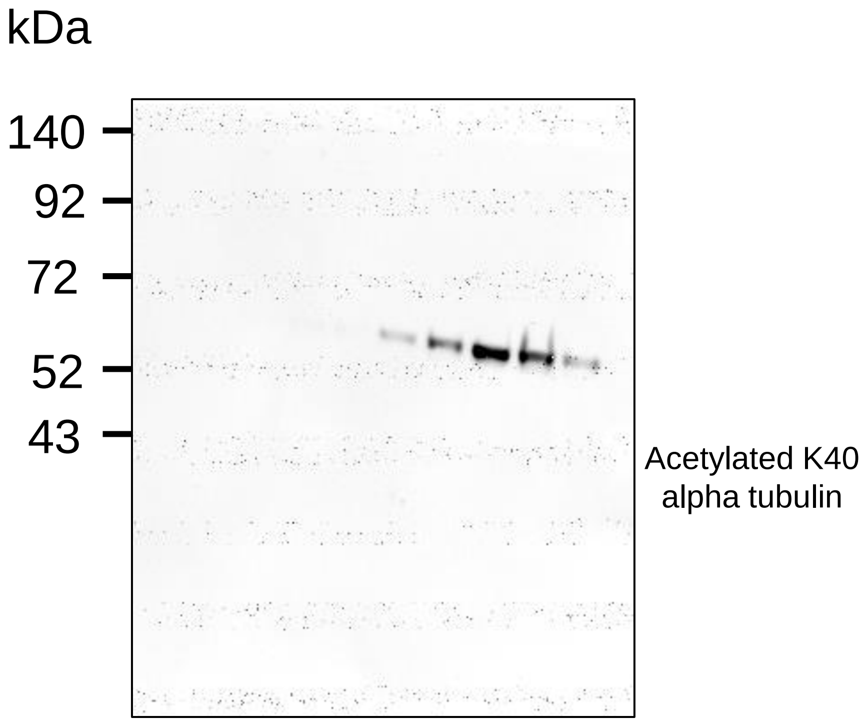


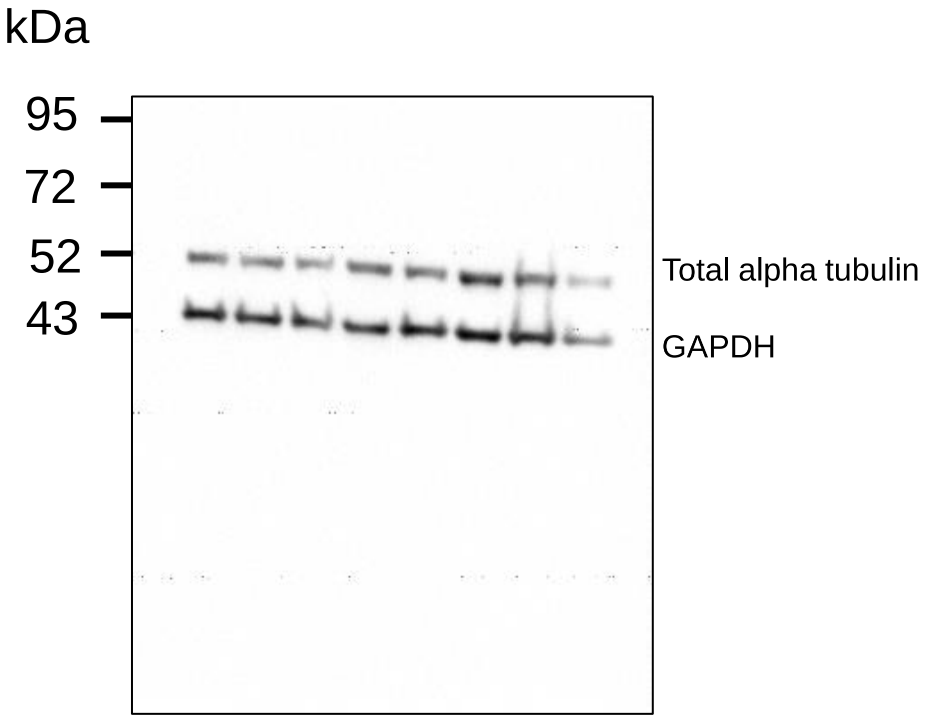

**Supplementary Figure S6C uncropped WB**. Some lanes removed as loaded with lysates not part of this study.

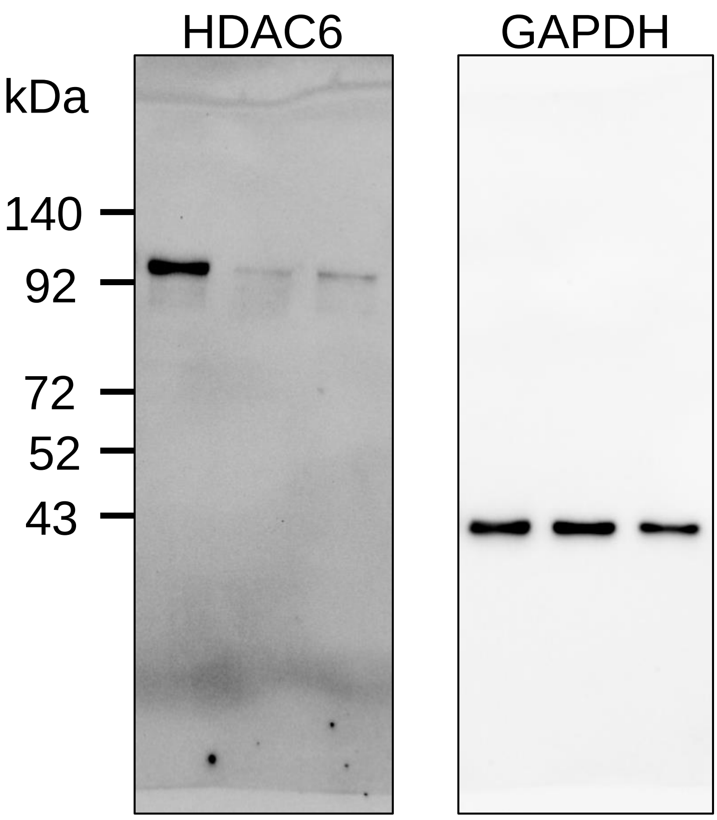

**Supplementary Figure 7A uncropped WBs.**

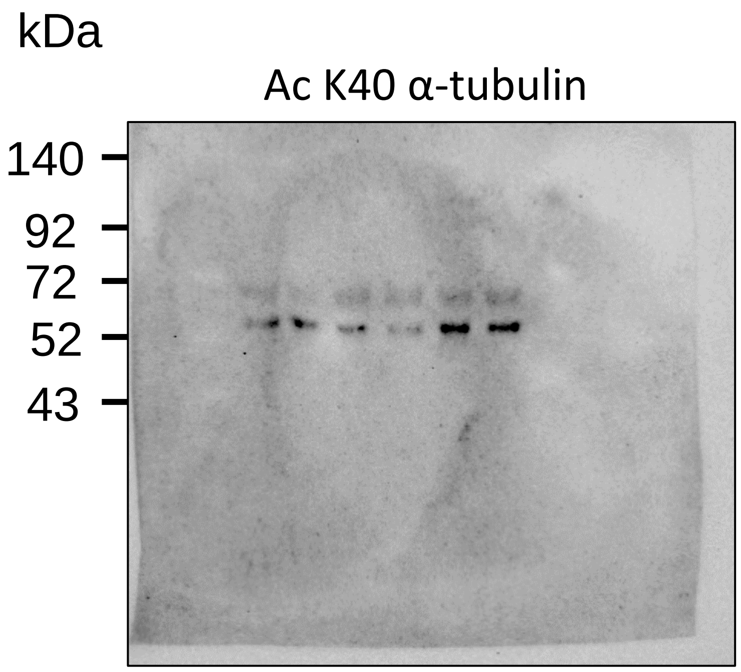


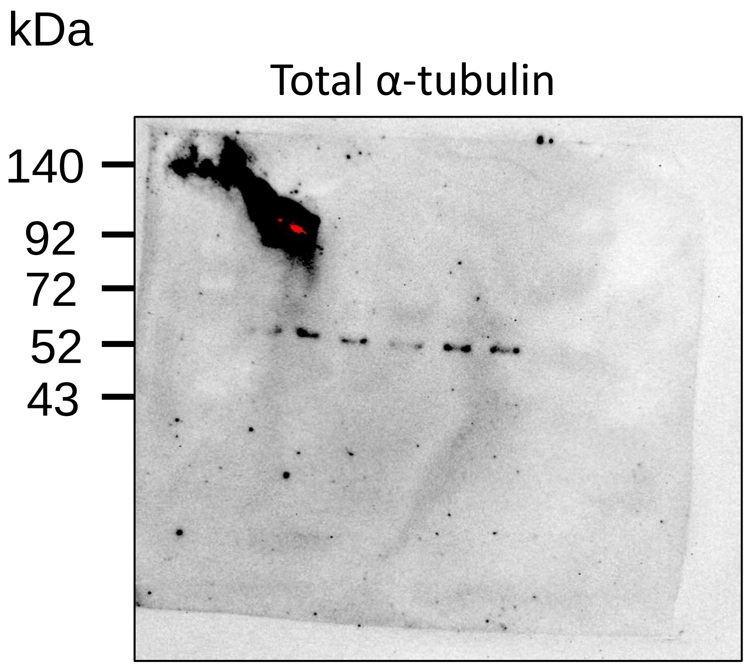


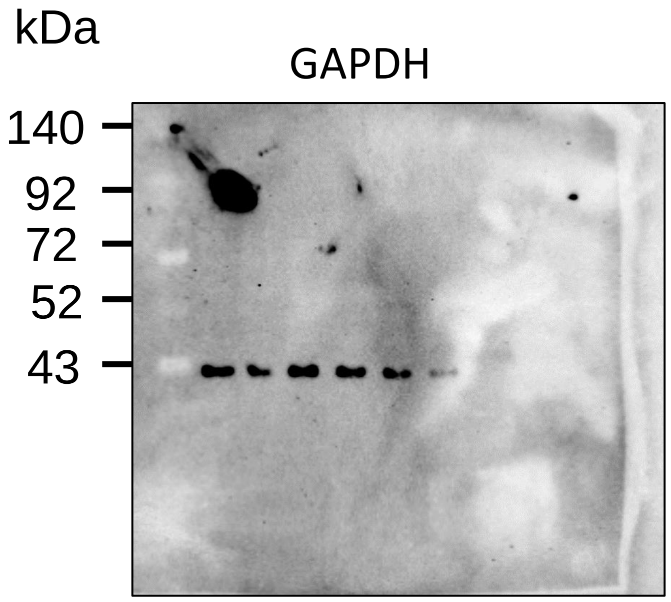
